# Antigen-driven PD-1^+^*TOX*^+^*EOMES*^+^ and PD-1^+^*TOX*^+^*BHLHE40*^+^ synovial T lymphocytes regulate chronic inflammation *in situ*

**DOI:** 10.1101/2019.12.27.884098

**Authors:** Patrick Maschmeyer, Gitta Anne Heinz, Christopher Mark Skopnik, Lisanne Lutter, Alessio Mazzoni, Frederik Heinrich, Sae Lim von Stuckrad, Lorenz Elias Wirth, Cam Loan Tran, René Riedel, Katrin Lehmann, Imme Sakwa, Rolando Cimaz, Francesco Giudici, Marcus Alexander Mall, Philipp Enghard, Bas Vastert, Hyun-Dong Chang, Pawel Durek, Francesco Annunziato, Femke van Wijk, Andreas Radbruch, Tilmann Kallinich, Mir-Farzin Mashreghi

## Abstract

T lymphocytes accumulate in inflamed tissues of patients with chronic inflammatory diseases (CIDs) and express pro-inflammatory cytokines upon re-stimulation *in vitro*^1–29^. Further, a significant genetic linkage to MHC genes suggests that T lymphocytes play an important role in the pathogenesis of CIDs including juvenile idiopathic arthritis (JIA)^30–33^. However, the functions of T lymphocytes in established disease remain elusive. Here we dissect the heterogeneity of synovial T lymphocytes in JIA patients by single cell RNA-sequencing. We identify subpopulations of T lymphocytes expressing genes reflecting recent activation by antigen *in situ*. A PD-1^+^*TOX*^+^*EOMES*^+^ population of CD4^+^ T lymphocytes expressed immune regulatory genes and chemoattractant genes for myeloid cells. A PD-1^+^*TOX*^+^*BHLHE40*^+^ population of CD4^+^, and a mirror population of CD8^+^ T lymphocytes expressed genes driving inflammation, and genes supporting B lymphocyte activation. This analysis points out that multiple types of T lymphocytes have to be targeted for therapeutic regeneration of tolerance in arthritis.

## Main Text

Here we have analyzed CD4^+^CD45RO^+^CD25^−^, CD4^+^CD45RO^+^CD127^lo^CD25^+^, and CD8^+^CD45RO^+^ T lymphocytes, representing antigen-experienced conventional CD4^+^ T lymphocytes (Tcon), regulatory CD4^+^ T lymphocytes (Treg) and CD8^+^ T lymphocytes, respectively. Cells were isolated from synovial fluid (SF) and peripheral blood (PB) of 7 patients with oligoarticular JIA by fluorescence-activated cell sorting (FACS), and single-cell transcriptomes and T cell receptor repertoire were determined by RNA-sequencing (Extended Data Fig. 1a and b; Supplementary Table 1). We had shown before that the isolation procedure used here does not affect gene expression of the isolated T cells, thus, these transcriptomes reflect gene expression *in vivo*^34–36^. We then performed shared nearest neighbor-clustering based on the transcriptional profiles of these cells and projected the resulting clusters using dimensional reduction analysis by t-distributed stochastic neighbor embedding (t-SNE)^37^. Most of the 74,891 cells of blood and synovia from all patients clustered according to their lineage and tissue origin (Fig. 1a).

**Figure 1:**
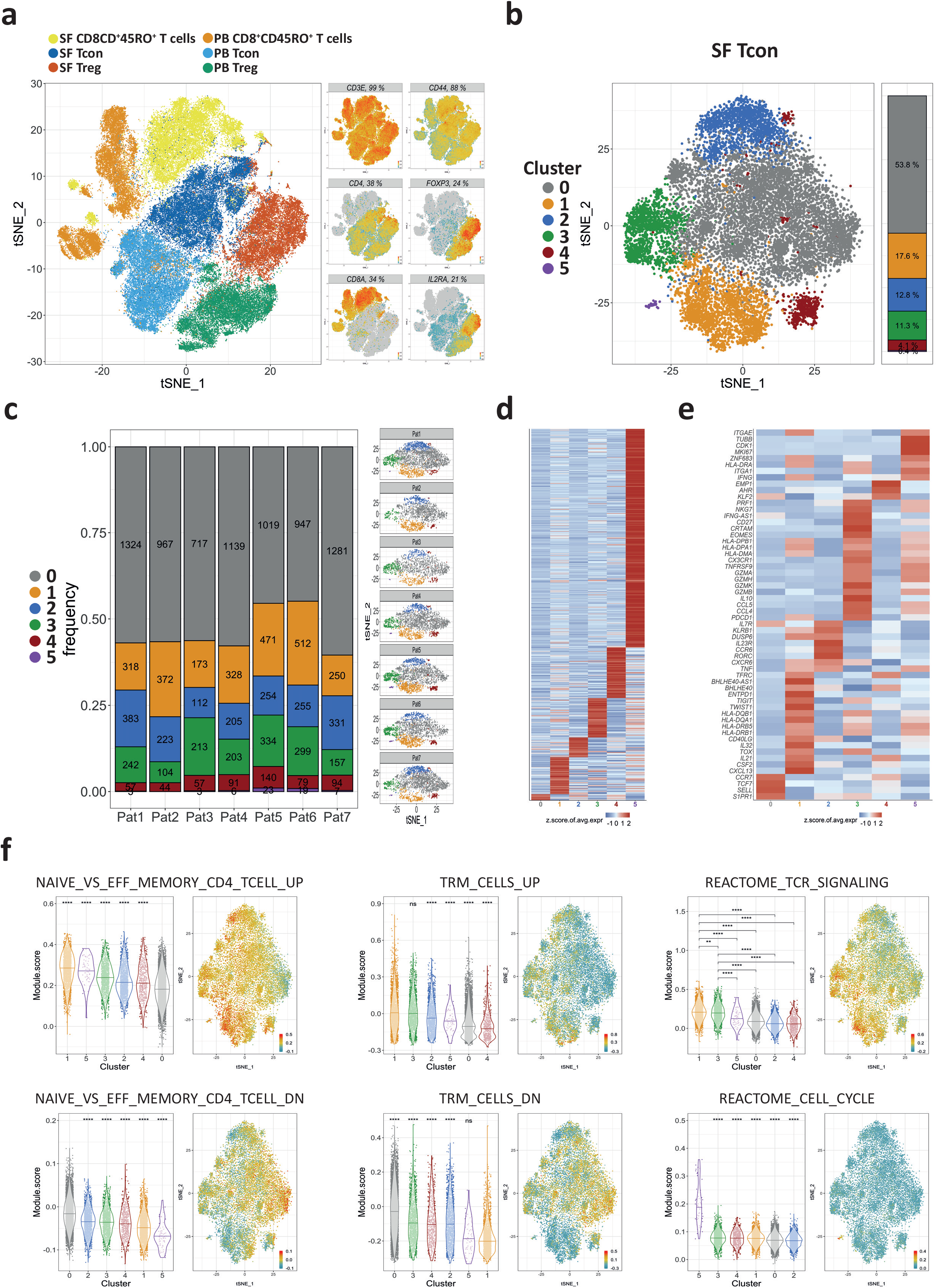
Transcriptional landscape of the antigen-experienced (memory) T cell compartment and heterogeneity of synovial CD4^+^ Tcon in inflamed joints of JIA patients. **a**, A t-SNE map of antigen-experienced T cells from the peripheral blood (PB) and the synovial fluid (SF) of JIA patients. The cells were FACS-sorted prior to single-cell RNA sequencing to determine the transcriptomes and the TCR repertoire. Shown is a t-SNE map of 74,891 CD4^+^CD45RO^+^CD25^−^ (Tcon), CD4^+^CD45RO^+^CD25^+^ (Treg) and CD8^+^CD45RO^+^ T cells from all patients combined. (n=7 for SF Tcon and SF CD8^+^CD45RO^+^ T cells; n=6 for SF Treg, PB Treg, PB Tcon and PB CD8^+^CD45RO^+^ T cells.). Each dot represents a single cell and is coloured according to the respective sorted population. **b**, CD4^+^ Tcon (n=13,756) from inflamed joints of 7 JIA patients were clustered and projected on a t-SNE map showing the formation of 6 different clusters (0-5). Each dot corresponds to a single cell and is colored according to its cluster affiliation. **c**, Quantification and t-SNE projection of the cluster distribution among synovial Tcon of the 7 different JIA patients. **d**, Heatmap showing the z-score normalized mean expression of 603 cluster-specific genes. **e**, Heatmap showing the z-score normalized mean expression of selected T cell function-associated genes. **f**, Module scores of gene sets associated to previously described T cell functions projected on t-SNE maps and statistical comparison between clusters illustrated as violin plots. In **d** and **e**, Heatmap color scales indicate row-wise z-scores of cluster-means of log-normalized expression values. Signature genes were tested for significant differential expression by cells of a cluster in comparison to cells from all other clusters by Wilcoxon’s rank sum test with Bonferroni correction. Cutoff was a minimum of 5 % expressing cells in the respective cluster and an adjusted p-value ≤ 0.05. In **f**, A two-sided unpaired Wilcoxon test with ns p > 0.05; *p ≤ 0.05; **p ≤ 0.01; ***p ≤ 0.001 and ****p ≤ 0.0001 was used. The gene set NAIVE_VS_EFF_MEMORY_CD4_TCELL_UP (GSE11057) contains genes, whose expression have been found to be upregulated in naive CD4^+^ T cells as opposed to effector memory T cells. The gene set NAIVE_VS_EFF_MEMORY_CD4_TCELL_DN (GSE11057) contains genes, whose expression have been found to be downregulated in naive CD4^+^ T cells as opposed to effector memory T cells. Analogously TRM_CELLS_UP and TRM_CELLS_DN, are gene sets containing genes that were found to be upregulated or downregulated in T_RM_ cells, respectively^52^. The curated REACTOME_TCR_SIGNALING (M15381) and REACTOME_CELL_CYCLE (M543) gene sets are from the Reactome database and contain genes that are associated with TCR signaling and with the cell cycle, respectively^51^. The particular genes of all used gene sets can be found in Supplementary Table 2.

Synovial Tcon from all patients combined were further clustered into six different subsets (clusters 0-5). All clusters were comprised of cells from each individual patient, except cluster 5, which did not include cells of patient 2 (Fig. 1b and c; Extended Data Fig. 2). Transcriptomic signatures for the 6 synovial Tcon clusters were defined by 603 genes (Fig. 1d). To characterize the clusters further, we analyzed their expression of functionally defined gene sets (modules) and individual signature genes (Fig. 1e and f, Supplementary Table 2)^38^. Cluster 0 was the largest, comprising an average of 53.8% (range: 44.9%-60.4%) of the synovial Tcon (Fig. 1c). Cells of cluster 0 expressed a gene module typical for naive or circulating central memory T (T_CM_) cells, including the signature genes *SELL, CCR7* and *S1PR1* (Fig. 1e and f; Extended Data Fig. 3)^39–45^. Cluster 1 comprised 17.6% (11.8%-24.25%) of the synovial Tcon (Fig. 1c). These cells expressed a gene module indicating activation by antigen in the synovia *in situ*, including MHC class II genes and *CD40LG*, as well as genes encoding for cytokines, like *IFNG, TNF, CXCL13* and *CSF2* (GM-CSF) (Fig. 1e and f; Fig. 3; Extended Data Fig. 3)^46–51^. These cells also expressed a gene module characterizing tissue-resident memory (T_RM_) cells, including genes such as *CXCR6* and *DUSP6*, *ITGAE* (CD103), *ZNF683* (HOBIT), and downregulation of *KLF2* (Fig. 1e and f; Extended Data Fig. 3)^45,52–55^. Cells of cluster 2 made up for 12.8% (8.8%-16.5%) of synovial Tcon (Fig. 1c). These cells qualified as *bona fide* T helper (T_h_) type 17 cells due to the expression of *RORC, IL23R*, and *CCR6* (Fig. 1e; Extended Data Fig. 3)^56^. A subset of these cells also expressed the T_RM_-associated gene module (Fig. 1f), and the genes *CD40LG, TNF* and *CSF2*, indicating their recent activation by antigen *in situ* (Fig. 1e; Fig. 3c)^45,47,52–55^. Cells of cluster 3 accounted for an average of 10% (6.1%-16.7%) of synovial Tcon (Fig. 1c). These cells expressed the gene module of recently activated T cells (Fig. 1f, REACTOME_TCR_SIGNALING), and genes indicating T cell receptor (TCR) stimulation, in particular *TNFRSF9* (4-1BB, CD137), CRTAM and MHC class II genes, but not *CD40LG* (Fig. 1e; Fig 3; Extended Data Fig. 3)^46,49–51,53,57–61^. They also expressed *EOMES* and/or *IL10* and/or *IFNG ex vivo*, resembling type 1 regulatory (Tr1)-like cells^62–64^. Cells of this cluster expressed gene modules of both circulating and T_RM_ cells (Fig. 1f), including the genes *S1PR1, CX3CR1* or *CXCR6* (Fig. 1e; Extended Data Fig. 3)^52–55^. Cells of cluster 4 represented 4.1% (2.5%-6.2%) of synovial Tcon (Fig. 1c). These cells expressed high levels of *CD44, ANXA1* (ANNEXIN A1), and/or *EZR* (EZRIN; Fig. 1e; Extended Fig. 3). Expression of these genes has been associated with survival, differentiation and functional polarization of T cells^65–69^. Some of the cells of this cluster also expressed *CD40LG* and cytokine genes, such as *CSF2* or *TNF* (Fig. 1e; Fig. 3). Cells of cluster 5 were a minor population (0.4%; 0.0%-1.0%) of all synovial Tcon (Fig. 1c). They were proliferating *in situ*, as indicated by expression of the Reactome gene module “cell cycle” (Fig. 1f), including genes like *MKI67* and *CDK1* (Fig. 1e)^51^. They also expressed MHC class II genes, indicating their recent activation by antigen (Fig 1e, Extended Data Fig. 3)^70–74^.

For 61.4% of the individual synovial Tcon (8445 cells), we could also determine the full-length TCR α and ß chain sequences. 5632 of these cells (66.7%) expressed a TCR detected only once, and 2813 cells (33.3%) expressed a TCR detected twice or more often (Fig. 2a). 512 (18.2%) of the detected expanded clones could be assigned to the 10 most abundant clonotypes, which possessed between 19 and 172 members (Fig. 2a). Cells of expanded clonotypes were most prevalent among cluster 1 (62.5%±14.2%), cluster 3 (41.7%±4.7%), and cluster 5, the latter in 3 out of 6 of the patients only (Fig. 2a and b).

**Figure 2:**
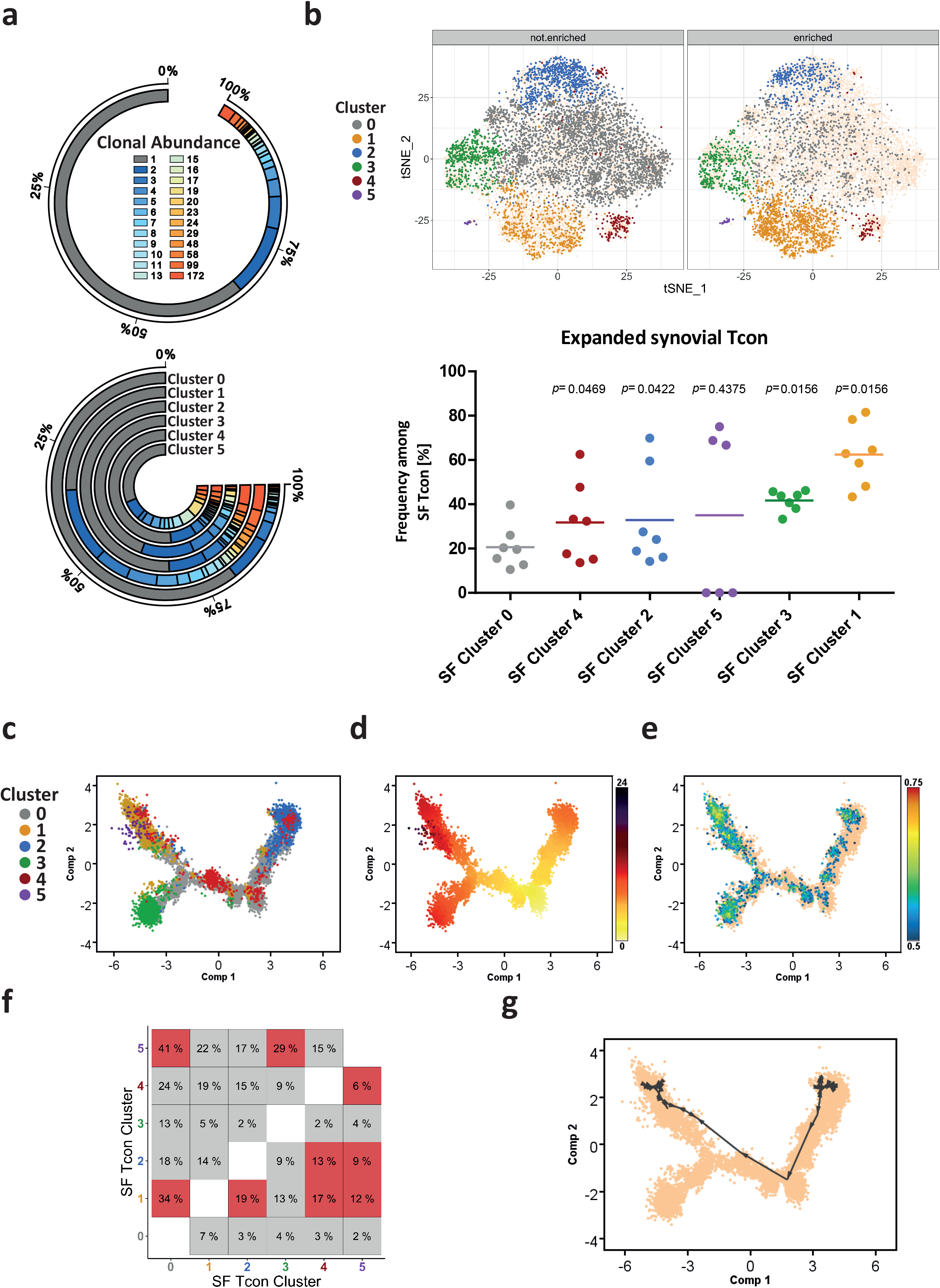
Clonal relationship of synovial Tcon reveals clusters 1 and 3 as clonally enriched and links cluster 1 to cluster 2 and 4. **a**, Detected frequencies of unique and expanded T cell clones among all synovial Tcon and in each cluster. The colour-scale corresponds to the clonal abundance (i.e. number of clones) of synovial Tcon clonotypes. **b**, Distribution of unique and expanded T cell clones among the t-SNE map of synovial Tcon as well as frequencies of expanded clones in each cluster. A two-sided paired Wilcoxon rank test was performed to determine significant differences between cluster 0 and each other cluster. Significance was assumed with a p ≤ 0.05. **c** and **d**, Pseudotime-based gene expression arranges cells along a Monocle trajectory resulting in a branched structure of synovial Tcon. **c**, Each cell is represented as one dot and labeled according to its affiliated cluster. **d**, Each cell is colored according to its pseudotime value, which suggests a direction of Tcon development with a root in the lower right corner (dominated by cluster 0). **e**, Projection of the REACTOME_TCR_SIGNALING (M15381) gene set on the trajectory. **f**, Clonal overlaps between the different clusters of synovial Tcon (in percent). Highlighted in red are clonal overlaps that are significantly overrepresented compared to 1000 simulated randomized overlaps. **g**, Shortest-Path analysis directed by clonal densities of Tcon clones that are present in all synovial clusters among the Monocle trajectory described in **c**. In b, **e** and **g**, Cells that are not highlighted are depicted as yellow in the background to outline the shape of the t-SNE map and the trajectory, respectively.

To define the relation between different subpopulations of synovial Tcon, we compared their TCR repertoires and generated a transcriptional trajectory according to gene expression along a pseudotime dimension with the unsupervised inference method Monocle, which organizes trajectories according to quantitative expression of signature genes^75^. We then projected the synovial Tcon clusters onto the trajectory (Fig. 2c and d, Extended Data Fig. 4 a). The resulting trajectory had a branched structure with 4 terminal nodes separated by two main components, with component 1 (x-axis) being determined by TCR signaling (Fig. 2e) and component 2 (y-axis) being dominated by signature genes of the distal nodes, which were defined by clusters 0, 1, 2 and 3 of synovial Tcon (Fig. 2c, Extended Data Fig. 4 a).

Comparison of the TCR repertoires of the synovial Tcon clusters showed overlaps between cells of clusters 1 and 0 (34%), 1 and 2 (19%), 1 and 4 (17%), 1 and 5 (12%), 2 and 4 (13%), 2 and 5 (9%), 4 and 5 (6%), 5 and 0 (41%), as well as cells of clusters 5 and 3 (29%). These overlaps were significantly overrepresented compared to simulated randomized distributions of TCR sequences among the different clusters (Fig. 2f, Extended Data Fig. 4b, Supplementary Table 3). Integrating the repertoire relationships and the trajectory based on expression of functional genes, a significant linkage of clusters 2, 4 and 1 became apparent. Shortest-Path analysis of the data suggested that T_h_17-like cells of cluster 2 had differentiated via a transitional state, represented by cluster 4, into antigen activated Tcon of cluster 1 (Fig. 2g), although cluster 1 also contained cells originating from clusters other than 2 and 4 (Extended Data Fig. 4b). In summary, clonal expansion, clonal linkage between clusters and a trajectory that is based on expression of functional genes, identify cells of clusters 1 and 3 as terminally differentiated, *in situ* activated Tcon putatively driving chronic inflammation of the joint.

Few of the cells of clusters 1 and 3 expressed *MKI67* or *CDK1*, suggesting that most of them were no longer proliferating, and had been clonally expanded in the past (Fig. 1e and f, Extended Data Fig. 3). Apart from the observed clonal expansion of Tcon from synovial clusters 1 and 3, those cells also expressed *TOX*, a gene indicating a history of repeated antigenic restimulation (Fig. 3b). Expression of *TOX* has been described to be induced by TCR stimulation in CD8^+^ T cells of a chronic infection mouse model, facilitating the survival of these cells^76–78^. The present analysis identifies expression of *TOX* as a hallmark of distinct CD4^+^ T cells of chronically inflamed human tissue for the first time. Among the genes known to be induced by *TOX* are *TIGIT* and *PDCD1*, which were also expressed by cells of clusters 1 and 3 (Fig. 3b)^78^. This phenotype of “exhaustion” contrasts with their expression of genes encoding cytokines and chemokines (Fig. 3c). It is intriguing that the cells expressing *TOX* or *PDCD1* are the cells expressing functional genes encoding cytokines and chemokines (Extended Data Fig. 5a and b). Quantitative PCR (qPCR) of flow-sorted synovial Tcon of JIA patients further confirmed that cells expressing the protein PD-1 also expressed higher levels of genes encoding for cytokines and chemokines than PD-1^−^ Tcon *in situ* (Extended Data Fig. 5c). Synovial PD-1-expressing Tcon also expressed higher levels of *TWIST1* and *EOMES*, genes encoding for transcription factors that regulate adaptation of T cells to chronic inflammation and control the expression of cytotoxicity-associated molecules, respectively^23,79,80^. Of note, cells of both clusters 1 and 3 lacked *CXCR5* expression, resembling non-follicular peripheral T_h_ cells that have been nominated as prime candidates driving chronic inflammation in rheumatoid arthritis and celiac disease (Fig. 3a, Extended Data Fig. 7b)^17,81^.

**Figure 3:**
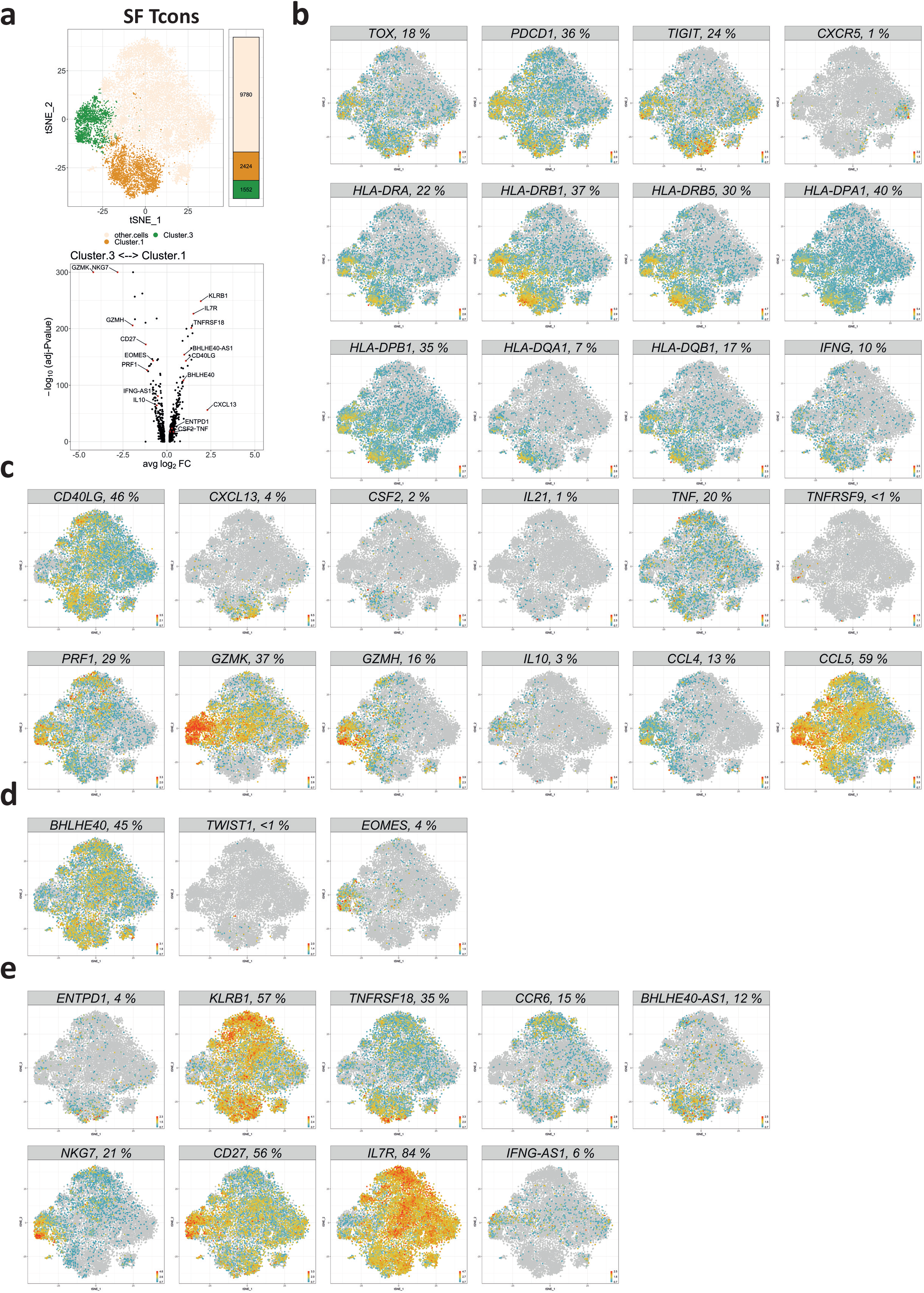
Subsets of antigen-stimulated *PDCD1*^hi^*TOX*^+^ synovial Tcon from JIA patients share a signature of activation but are distinct regarding their functional responses. **a**, A t-SNE map highlighting synovial Tcon in clusters 1 and 3 and a volcano plot showing genes that are differentially expressed by cells in these clusters. Significance was determined by Wilcoxon’s rank sum test with Bonferroni correction. Cutoff was a minimum of 5% expressing cells in either cluster. **b**, Feature plots showing the expression of indicated genes whose expression (or absence thereof, e.g. CXCR5) are shared between cells of cluster 1 and 3. **c**, Feature plots of indicated functional response genes that are differentially expressed between synovial Tcon of clusters 1 and 3. **d**, Feature plots of indicated transcription factor genes that are differentially expressed between cells of synovial Tcon clusters 1 and 3. **e**, Expression of indicated marker genes to distinguish between cells of clusters 1 and 3 among synovial Tcon. In **b-d**, Percentages in header bars of feature plots display the frequencies of Tcon that express the indicated genes.

Despite their common expression of *TOX, PDCD1* and of other genes indicating recent cognate activation, cells of cluster 1 and cluster 3 differed fundamentally in their response to antigenic stimulation (Fig. 3a-c). Cells of cluster 1 expressed *CD40LG* and pro-inflammatory cytokine and chemokine genes, in particular *CSF2, TNF, IFNG, IL21* and *CXCL13* (Fig. 3a and c; Extended Data Fig. 5d). Some of these cells also expressed CD25 as a hallmark of recent activation *in situ*, since CD4^+^CD45RO^+^CD25^+^ T cells from the same synovia identified a cluster of FOXP3^−^ T cells that resembled CD25^−^ T cells of synovial cluster 1 with respect to expression of cytokines and chemokines (see cluster 2 of CD4^+^CD45RO^+^CD127^lo^CD25^+^ T cells in Extended Data Fig. 6).

Tcon of synovial cluster 3 responded differently to antigenic stimulation *in situ* than Tcon of synovial cluster 1. They expressed *TNFRSF9* (4-1BB, CD137), *IL10, PRF1* and genes encoding granzymes (Fig. 3a and c), i.e. genes associated with Tr1-like cells. Such cells have been reported to regulate immune reactions by IL-10, and to kill myeloid cells and activated T cells with granzymes^62–64,82–85^. Nevertheless, they may also support inflammation, since they expressed the chemokine genes *CCL4* and *CCL5* (Fig. 3c), with the potential to attract myeloid cells^83,86–88^. Interestingly, some cells of synovial Tcon cluster 1 expressed *CSF2* and *TNF*, indicating that they may support activation of myeloid cells, while others expressed *IL21* and *CXCL13*, indicating that they have the potential to attract and support B cell activation and differentiation (Fig. 3c)^89–96^.

Cells of clusters 1 and 3 also differed fundamentally in their expression of key transcription factors. While cells of cluster 1 expressed *BHLHE40* and *TWIST1*, cells of cluster 3 expressed *EOMES* (Fig. 3d, Extended Data Fig. 7a). *TWIST1* is selectively upregulated in CD4^+^ T cells upon repeated antigenic stimulation and enhances expression of genes promoting persistence of the cells^25,26,97^. 34 of 41 synovial Tcon that expressed *TWIST1*, belonged to cluster 1 (Fig. 3d). While *TWIST1*^+^ cells were sparsely detectable by single-cell sequencing, *TWIST1* among synovial Tcon cells of cluster 1 was readily detectable by qPCR (Extended Data Fig. 7a). Expression levels of *TWIST1* in T cells are dependent on antigen-receptor activation and they decline rapidly, already 3 hours after onset of activation, highlighting cells that had been stimulated by antigen within 3 hours before biopsy^23^. Cells of cluster 1 also expressed the gene encoding BHLHE40 (Fig. 3a and d). This transcription factor has been described to induce the expression of *CSF2*, and to suppress the expression of *IL10^98–100^*. This is in line with the expression pattern we observe here for synovial Tcon of cluster 1. Finally, in line with the functional genes expressed by Tr1-like Tcon of synovial cluster 3 (Fig. 3a-c; Extended Data Fig. 5a and b), these cells expressed the transcription factor *EOMES*, which has been reported to control the differentiation of Tr1-like cells (Fig. 3d)^62,63^.

Apart from the genes discussed so far, Tcon of synovial clusters 1 and 3 differentially expressed other genes. Cells of cluster 1 expressed *ENTPD1* (CD39), *KLRB1* (CD161), *CCR6*, *TNFRSF18* (GITR) and the long-non-coding RNA *BHLHE40-AS1* (Fig. 3e). Cells of cluster 3 did not express *IL7R*, but *CD27*, *NKG7* and the long-non-coding RNA *IFNG-AS1* (Fig. 3e). These genes may serve in the future to identify and characterize such cells in pathological context.

PD-1 and CD39 and/or CD161 are sufficient to identify and discriminate synovial CD4^+^ T cells of clusters 1 and 3 in JIA (Extended Data Fig. 7). Comparing expression of the genes *CXCL13, IL21, CSF2, TWIST1, IFNG, BHLHE40AS1, IL10* and *EOMES* of PD-1^+^CD39^−^ and PD-1^+^ CD39^+^ synovial CD4^+^ T cells confirmed the gene expression patterns observed by single cell RNA-sequencing. In addition, flow cytometric analysis showed that PD-1^high^CXCR5^−^CD4^+^ T cells accumulated in the inflamed joints of JIA patients and that the dichotomy of these cells remained stable after re-stimulation *in vitro*. Pro-inflammatory PD1^high^CD4^+^ T cells expressing CD39 and/or CD161 secreted the cytokines GM-CSF and IL-21, while EOMES^+^ Tr1-like cells secreted IL-10 (Extended Data Fig. 7b-e). Mutually exclusive expression of CD161 and CD39 versus EOMES among T_h_ cells, and the accumulation of CD161^+^CD4^+^ T cells was observed not only for inflamed joints of JIA, but also for inflamed intestines of patients with inflammatory bowel disease (IBD; i.e. Crohn’s disease and ulcerative colitis), suggesting that PD1^high^CD39^+^/CD161^+^ and PD1^high^CD39^−^/CD161^−^ CD4^+^ T cells are involved in chronic inflammation of various tissues in distinct diseases (Extended Data Fig. 8a-c).

While synovial Tcon of cluster 1 expressed only gene modules and signature genes of T_RM_ cells, synovial Tcon of cluster 3 showed expression of genes associated both with resident and circulating T cells (Fig. 1e and f, Extended Data Fig. 3). Accordingly, we could detect Tcon in the peripheral blood of JIA patients, resembling Tr1-like cells of synovial Tcon cluster 3. For this, we analyzed the clonality of CD4^+^ blood Tcon, which possessed a reduced abundancy of expanded clones compared to synovial CD4^+^ Tcon (Fig. 4d; Extended Data Fig. 9a). This analysis revealed that CD4^+^ Tcon of blood cluster 5 harbored the highest frequency of expanded CD4^+^ Tcon clones among all blood clusters, as well as clones that overlap with CD4 Tcon clones of paired synovial fluid samples (Fig. 4d; Extended Data Fig. 9b). Tcon of blood cluster 5 specifically recapitulated the antigen receptor repertoire of synovial Tcon cluster 3, in that 37% of clones were shared between cells of both clusters in the 6 patients from who we received both blood and synovial fluid (Fig. 4e, Supplementary Table 4). Accordingly, Tcon of blood cluster 5 localized next to Tcon of synovial cluster 3, in a combined t-SNE map (Fig 4f). Tcon of blood cluster 5 were highly similar to Tcon of synovial cluster 3 and expressed functional molecules of these Tr1-like cells (Fig. 4g). However, they did not express *IL10 in situ* (Extended Data Fig. 9c). This may indicate that the circulating cells had not recently been activated by (synovial) antigens or lacked recent exposure to additional factors specifically found in inflamed tissues. Tcon of blood cluster 5, with their Tr1-like transcriptome, might define a novel immunoregulatory mechanism preventing systemic spread of inflammation and confining inflammation to distinct joints in oligoarthritis.

**Figure 4:**
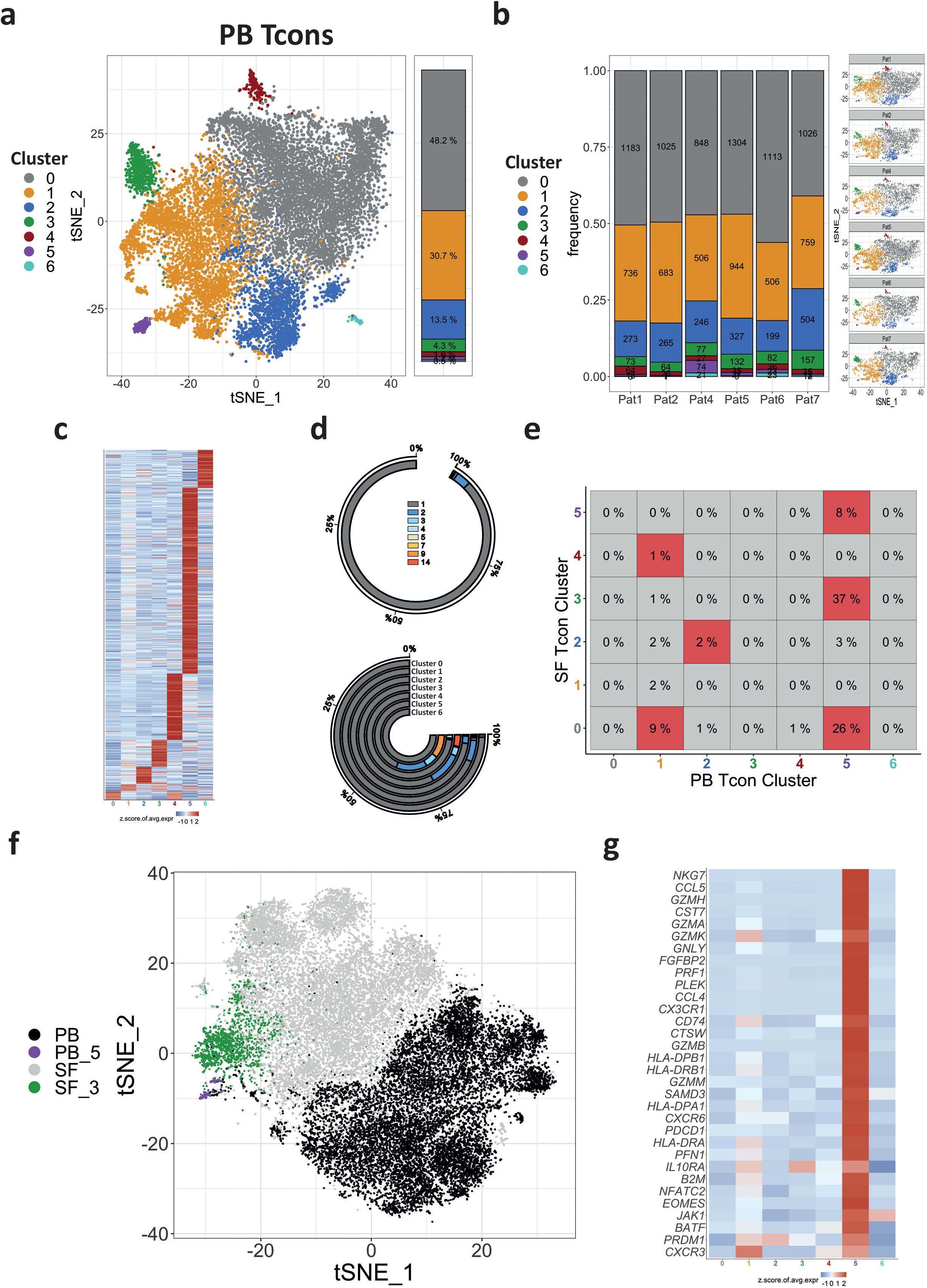
Tr1-like cells from synovial Tcon cluster 3 of inflamed joints are circulating in the peripheral blood of JIA patients. **a**, Cluster analysis and t-SNE projection of 13,479 Tcon from the PB of 6 JIA patients. Each dot represents a single cell and is colored according to its affiliation to one of the 7 identified clusters (clusters 0-6). **b**, Quantification and t-SNE projections of clusters among the individual JIA patients. **c**, Heatmap showing the z-score normalized mean expression of 402 cluster-specific genes. **d**, Detected frequencies of unique and expanded T cell clones among all circulating PB Tcon and in each PB Tcon cluster. The color-scale corresponds to the clonal abundance (i.e. number of clones) of PB Tcon clonotypes. **e**, Clonal overlaps between the different clusters of synovial and PB Tcon (in percent). Highlighted in red are clonal overlaps that are significantly overrepresented compared to 1000 simulated randomized overlaps. **f**, Combined t-SNE map of Tcon from the PB and the SF. Each dot represents a cell. Displayed are cells from PB Tcon cluster 5 (violet), PB Tcon from all clusters except cluster 5 (black), SF cluster 3 (green) and Tcon from all SF clusters except from cluster 3 (grey). **g**, Heatmap displaying genes that are expressed by circulating Tr1-like cells found in PB cluster 5 representing a molecular fingerprint for the identification of SF derived cells in the PB. In **c** and **g**, Heatmap color scales indicate row-wise z-scores of cluster-means of log-normalized expression values. Signature genes were tested for significant differential expression by cells of a cluster in comparison to cells from all the other clusters by Wilcoxon’s rank sum test with Bonferroni correction. Cutoff was a minimum of 5 % expressing cells in the respective cluster and an adjusted p-value ≤ 0.05.

Apart from Tcon, also CD8^+^ T memory cells have been implicated earlier in the pathogenesis of JIA^29^. Here we originally identified a prominent cluster of synovial CD8^+^ T cells with transcriptomes resembling those of pro-inflammatory synovial CD4^+^ T helper cells of cluster 1. Synovial CD8^+^CD45RO^+^ T lymphocytes of all 7 JIA patients robustly and reproducibly clustered into 10 different populations (Fig. 5a). Their frequencies were between 0.1% and 57.6% of the 14,019 synovial CD8^+^CD45RO^+^ T lymphocytes as indicated in Fig. 5b, and differential gene expression signatures defined by 851 genes (Fig. 5c). Cells of most clusters expressed *PDCD1, TOX* and *IFNG*, indicating their cognate activation *in situ*. Synovial CD8^+^CD45RO^+^ T cells of cluster 1 represented 15.6% (11.6%-16.4%) of synovial CD8^+^CD45RO^+^ T cells. These cells upregulated expression of the genes *CD40LG, CSF2, IL21, TNF, KLRB1* (CD161), *ENTPD1* (CD39), *TNFRSF18* (GITR), *CXCL13* and also the long-non-coding RNA *BHLHE40-AS1*. In parallel, they downregulated the expression of *EOMES, NKG7* or genes encoding granzymes as compared to cells outside of cluster 1. This was also true for the genes *CCR7, SELL, KLF2, S1PR1* and *CXCR5* (Fig 5d and e). Of note, PD-1^high^CD8^+^ T cells were enriched in inflamed joints of JIA patients compared to blood and were able to secrete also the proteins IL-21 and GM-CSF (Extended Data Fig. 10) and expressed both *CD8B* and *CD8A* and signature genes of T_RM_ cells, such as *ITGA1* (CD49b), *DUSP6* and *CXCR6* (Fig. 5e). Taken together, their distinct gene expression signature indicates that these cells *i)* are activated by antigen, and *ii)* are presumably an MHC class I-restricted CD8^+^ “mirror population” of pro-inflammatory synovialCD4^+^ Tcon of cluster 1, and thus, cells with the potential to drive chronic inflammation^29^.

**Figure 5:**
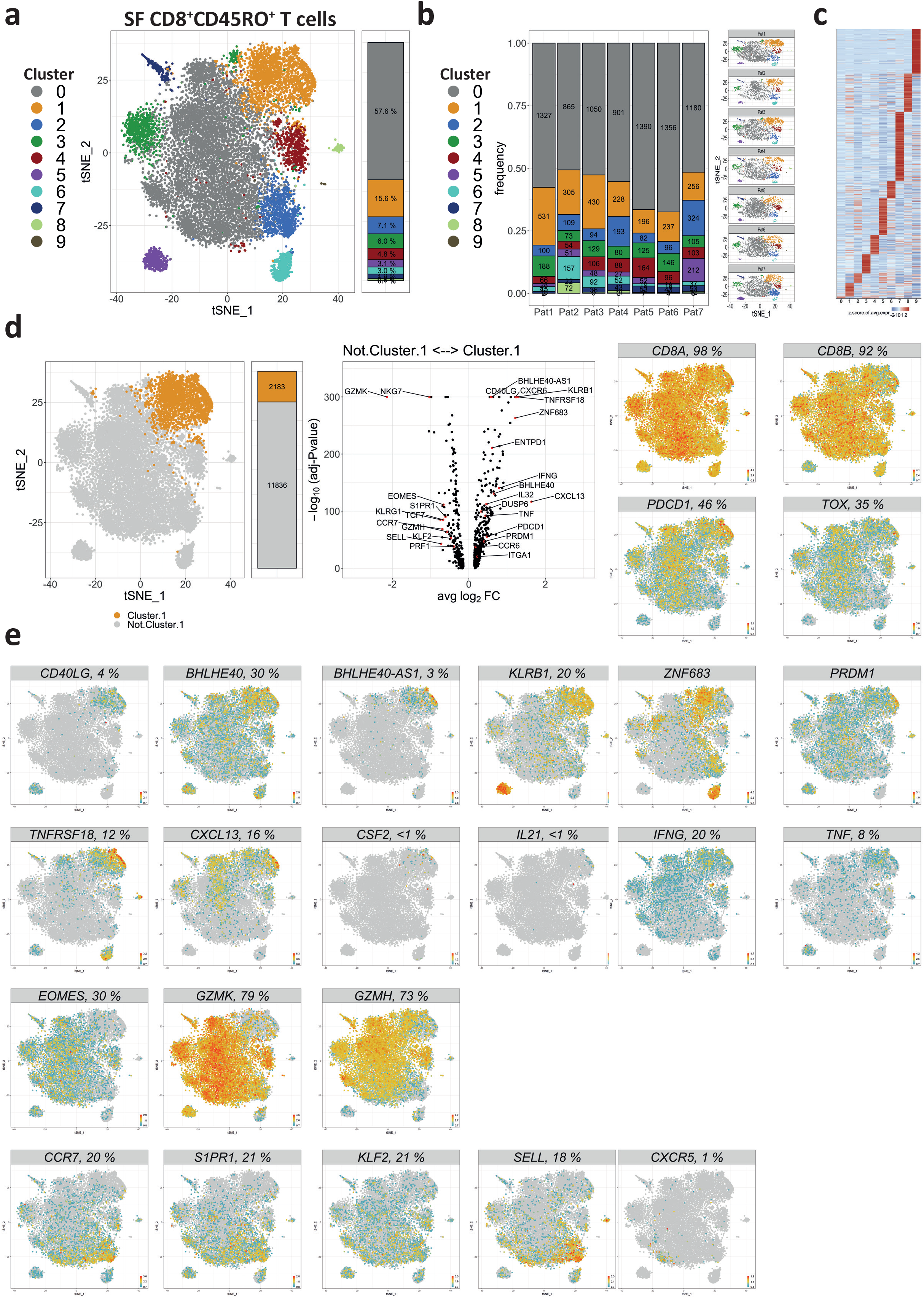
A subset of antigen-experienced (memory) CD8^+^ T cells resembles pro-inflammatory T helper cells in the inflamed joints of JIA patients and expresses a T_RM_ cell gene signature. **a**, 14,019 CD8^+^CD45RO^+^ T cells from inflamed joints of 7 JIA patients were sequenced, clustered and projected onto a t-SNE map. Clustering separated them into 10 different subsets (clusters 0-9). Each dot represents a single cell and is colored according to its cluster affiliation. **b**, Quantification and t-SNE projections of the cluster distributions among individual patients. **c**, Heatmap showing the z-score normalized mean expression of 851 cluster-specific genes. **d**, Volcano plot summarizing differentially expressed genes of CD8^+^CD45RO^+^ T cells in synovial cluster 1, as compared to synovial CD8^+^CD45RO^+^ T cells that are not present in cluster 1. In addition, a t-SNE map showing the groups with the cells that were compared and feature plots depicting the expression of *CD8A, CD8B, PDCD1* and *TOX* are shown. Significance was determined by Wilcoxon’s rank sum test with Bonferroni correction. Cutoff was a minimum of 5% expressing cells in either cluster. **e**, Feature plots showing the expression of indicated genes whose expression is up- or downregulated by cells of SF CD8^+^CD45RO^+^ T cells in cluster 1.

In summary, dissecting synovial effector T lymphocytes from patients with JIA using single-cell transcriptomics revealed an unforeseen heterogeneity and plasticity. Many of the cells express signature genes indicating continued activation by antigen *in situ* and transcribe genes encoding cytokines and chemokines with a role in inflammation. Although clusters of T cells can be defined according to their transcriptional similarity, individual reaction patterns to antigenic stimulation differ, even within clusters. Of particular interest are a PD-1^+^*TOX*^+^*EOMES*^+^ population of CD4^+^ T lymphocytes, representing clonally expanded, probably terminally differentiated, non-proliferating but very active effector T cells, potentially attracting myeloid cells, but also expressing *IL10* with the potential to regulate the immune reaction. These cells are also found in the circulation and might have the potential to limit systemic spread of inflammation. In any case, they may serve as biomarkers to monitor JIA disease activity and response to therapy non-invasively. Two other populations of interest are a PD-1^+^*TOX*^+^*BHLHE40*^+^ population of CD4^+^, and a mirror population of CD8^+^ T lymphocytes, presumably supporting extra follicular B cell activation by secretion of IL-21 and CXCL13, as well as activation of myeloid cells by secretion of TNF and GM-CSF. These cells, too, are clonally expanded, non-proliferating and terminally differentiated. The considerable individual heterogeneity of cells driving inflammation in various ways, and limiting it in others, provides a challenge for the development of effective targeted therapies that address these diverse cell types differentially and concomitantly, to be efficient. The present data provide a molecular basis for the development of biomarkers and targeted immune modulating therapies for JIA and potentially other chronic inflammatory diseases.

## Materials and Methods

### Human patient samples

Peripheral blood (PB) and synovial fluid (SF) samples of JIA patients were collected at the Department of Pediatric Pulmonology, Immunology and Critical Care Medicine, Division of Pediatric Rheumatology at Charité–Universitätsmedizin Berlin as approved by the ethics committee of the Charité–Universitätsmedizin Berlin (approval no. EA2/069/15), at the Pediatric Rheumatology Department at University Medical Center of Utrecht (The Netherlands) in accordance with the Institutional Review Board of the University Medical Center Utrecht (approval no. 11-499/C), and at the Meyer children’s hospital, Florence, as approved by the Meyer children’s hospital ethics committee (approval no. 184/2016). PB from healthy adult volunteers was obtained from the Mini Donor Service at University Medical Center Utrecht. Samples were collected in compliance with the Declaration of Helsinki. Peripheral blood (PB) and synovial fluid (SF) was obtained via vein puncture or intravenous drip, and by therapeutic joint aspiration of the affected joints, respectively. Informed consent was obtained from all patients either directly or from parents/guardians. Intestinal biopsies of IBD patients were obtained from the Azienda Ospedaliero-Universitaria Careggi (AOUC), Florence, as approved by the ethics committee of AOUC hospital (approval no. 12382_BIO).

### Isolation of antigen-experienced (memory) T cells from the SF and the PB of JIA patients for single-cell sequencing and qPCR analyses

Mononuclear cells from the peripheral blood (PBMCs) were isolated by Ficoll density-gradient centrifugation and subsequent collection of the leukocyte-containing interphase.

For the isolation of leukocytes from the synovial fluid, we diluted the viscous fluid with 5 mM ethylenediaminetetraacetic acid (EDTA)-containing phosphate-buffered saline (PBS) and filtered the cells using a 70 μm cell strainer (BD). Subsequently, cells were centrifuged at 300 g at 4 °C and resuspended in PBS containing 0.2% bovine serum albumin (PBS/BSA). Following an additional filtration step through a 70 μm cell strainer, cells were resuspended in PBS/BSA prior to depletion of CD15+ neutrophils by magnetic cell separation (MACS, Miltenyi Biotec).

Single cell suspensions of PBMCs and CD15-depleted SF cells were blocked with human FcR blocking reagent (Miltenyi Biotec) and labeled for 15 min at 4 °C with antibodies for fluorescent activated cell sorting (FACS) using a FACS-Aria II (BD). The following antibodies were used for labeling cells prior to FACS: anti-human CD3-Alexa Fluor 405 (AF405, clone UCHT1, DRFZ Berlin), CD4 Brilliant Violet 785 (BV785, EH12.2H7; Biolegend), CD45RO (BV510, clone UCHL1, BD), CD39 (BV605, clone A1, Biolegend), CD127 (AF488, clone A019D5, Biolegend), CD25 allophycocyanin (APC, M-A251, BD), CD8a APC-Cyanine 7 (APC-Cy7, clone: HIT8a, Biolegend), PD-1 phycoerythrin (PE)-Cy7 (clone EH12.2H7, Biolegend), CD14 peridinin-chlorophyll protein complex (PerCP, clone TM1, DRFZ Berlin) and propidium iodide (PI, Sigma Aldrich).

Cells were sorted to obtain antigen-experienced (memory) CD4^+^CD45RO^+^CD25^−^ T cells (Tcons), CD4^+^CD45RO^+^CD127^lo^CD25^+^ regulatory T cells (Treg), and CD8^+^CD45RO^+^ T cells as shown in Extended Data Fig. 1. Sorted cells then were subjected to single-cell sequencing or qPCR following isolation or RNA.

### Isolation of T cells from the SF and the PB for confirmation of single-cell sequencing data by flow cytometry

SF of JIA patients was incubated with hyaluronidase (Sigma-Aldrich) for 30 min at 37°C to break down hyaluronic acid. Synovial fluid mononuclear cells (SFMCs) and peripheral blood mononuclear cells (PBMCs) were isolated using Ficoll Isopaque density gradient centrifugation (GE Healthcare Bio-Sciences, AB) and frozen in Fetal Calf Serum (FCS) (Invitrogen) containing 10% DMSO (Sigma-Aldrich) until use.

To perform analysis of cells by flow cytometry, the cells were thawed and transferred to a 96-well plate and stained for viability with Viability Dye eFluor 506 (eBioscience) followed by incubation with the surface antibodies: anti-human CD3 AF700 (clone UCHT1, Biolegend), CD4 BV785 (clone OKT4, Biolegend), CD8α PerCP-Cy5.5 (clone SK1, BD), CD127 BV605 (clone A019D5, Sony Biotechnology), CD25 BV711 (clone 2A3, BD), CD45RO (clone UCHL1, BD), CD39 fluorescein isothiocyanate (FITC, clone A1, Serotec), CXCR5 PerCP-Cy5.5 (clone J252D4, AntiBodyChain), CD161 PE-Cy5 (clone DX12, BD), CD161 APC (clone DX12, BD), CD279 BV711 (clone EH12.1, BD), CD279 PE (clone EH12.1, BD). For intracellular staining cells were fixed and permeabilized using the Intracellular Fixation & Permeabilization Buffer Set (eBioscience) followed by incubation with: FOXP3 eF450 (clone PCH101, eBioscience), IL-21 PE (clone 3A3-N2.1, BD), IL-10 APC (clone JES3-19F1, Biolegend), GM-CSF PE-Dazzle594 (clone BVD2-21C11, Biolegend), IFN-γ PE-Cy7 (clone 4S.B3, BD). For intracellular cytokine production, cells were first stimulated for a total of 4 hours with phorbol 12-myristate 13-acetate (PMA, 10 ng/ml; MP Biomedicals) and ionomycin (1 μg/ml; Calbiochem). Golgi stop (1/1500; BD Biosciences) was added for the last 3 hours of stimulation. Hereafter, the normal staining procedure was followed. Data acquisition was performed on a BD LSRFortessa (BD Biosciences) and analyzed using FlowJo Software (Tree Star Inc.).

Statistical analysis of cytokine expression (protein data) was performed with a one-way ANOVA with Dunnett post-hoc test and with a paired Student’s t-test, if applicable using GraphPad Prism version 7.04 for Windows (GraphPad Software, La Jolla California USA).

### Isolation and staining of T cells from IBD patients

Mononuclear cells from the peripheral blood (PBMCs) were isolated by Ficoll density-gradient centrifugation and subsequent collection of the leukocyte-containing interphase.

Biopsies of inflamed gut tissue were aseptically obtained during curatively-intended surgical procedures. Fresh samples were then minced with a scalpel and enzymatically digested for 90 min with Collagenase A (1 mg/ml) at 37°C on continuous rotation using MACS Mix (Miltenyi). Digested materials were then filtered with a 40 μm cell strainer and pelleted. Red cell lysis was eventually performed using 0.8% NH_4_Cl solution. Recovered cells from the PB and from gut biopsies were activated *in vitro* in RPMI-1640 medium + 10% FCS with PMA (10 ng/ml) and ionomycin (1 μM) for 5 hours, the last 3 hours in the presence of Brefeldin A (5 μg/ml). Subsequently, cells were washed with PBS and then fixed using the Transcription factor Staining Buffer Set from eBioscience, following the manufacturer’s instructions. Cells were then stained with anti-human CD3 Pacific Blue (clone UCHT1, BD), CD4 PE (SK3, BD), CD161 PE-Vio770 (191B8, Miltenyi Biotech), *EOMES* APC (WD1928, eBioscience). Finally, samples were acquired on a BD LSR II flow cytometer.

### RNA isolation and qPCR of cDNA

In order to determine gene expression by qPCR, sorted cells were lysed in Qiazol and stored at −80 °C. RNA was isolated from the lysate by using the miRNeasy Micro Kit (Qiagen) according to manufacturer’s instructions. Concentrations and purity of isolated RNA was measured with a Nanodop 2000. Reverse transcription was performed using the Reverse Transcription kit from Applied Biosystems. Gene expression was quantified with qRT-PCR based TaqMan Assays (ThermoFisher) using the following assays: *BHLHE40-AS1*: Hs04407259_m1, *CSF2*: Hs00929873_m1, *CXCL13*: Hs00757930_m1, *ENTPD1*: Hs00969556_m1, *EOMES*: Hs00172872_m1, *HPRT1*: Hs02800695_m1, *IFNG*: Hs00989291_m1, *IL10*: Hs00961622_m1, *IL21*: Hs00222327_m1, *TWIST1*: Hs01675818_s1,. Expression of genes was quantified with the (2^−ΔCT^) method relative to the expression of *HPRT1*.

### Single-cell RNA-sequencing

For single cell library preparation, FACS-sorted human T cells were applied to the 10X Genomics platform using the Chromium Single Cell 5’ Library & Gel Bead Kit (10x Genomics) and following the manufacturer’s instructions for capturing ~3000 cells. The amplified cDNA was used for simultaneous 5’ gene expression (GEX) and TCR library preparation. TCR target enrichment was achieved using the Chromium Single Cell V(D)J Enrichment Kit for Human T cells. Through fragmentation, adapter ligation and index PCR the final libraries were obtained. The quality of single cell 5’ GEX and TCR libraries was assessed by Qubit quantification, Bioanalyzer fragment analysis (HS DNA Kit, Agilent) and KAPA library quantification qPCR (Roche). The sequencing was performed on a NextSeq500 device (Illumina) using High Output v2 Kits (150 cycles) with the recommended sequencing conditions for 5’ GEX libraries (read1: 26nt, read2: 98nt, index1: 8nt, index2: n.a.) and Mid Output v2 Kits (300 cycles) for TCR libraries (read1: 150nt, read2: 150nt, index1: 8nt, index2: n.a., 20% PhiX spike-in). Raw Illumina-NextSeq 500 data were processed using cellranger-2.1.1. Mkfastq, count commands and vdj commands were used with default parameter settings. The genome reference was refdata-cellranger-hg19-1.2.0. The vdj-reference was refdata-cellranger-vdj-GRCh38-alts-ensembl-2.0.0. In both cases, the number of expected cells was set to 3000. Single-cell transcriptome and immune profiling data discussed in this publication, are available at gene expression omnibus (GEO) under the accession number XXX.

### Quality control and filtering of single-cell transcriptomes

Single cell RNA sequencing data were mapped as described above. The resulting expression matrix was analyzed in R using the Seurat package (version 3.1.1) following the principle steps described previously by Tallulah S. Andrews and Martin Hemberg^37^. In brief, putative artifacts from each sample were filtered by removing transcriptomes with less than 2,500 and more than 12,500 transcripts and transcriptomes comprising less than 1% or more than 10% of mitochondrial transcripts. In addition, also the 1^st^ and 95^th^ percentile of transcriptomes, as judged by the fraction of mitochondrial transcripts, as well as the 5^th^ and 95^th^ percentile, as judged by the number of transcripts, were discarded. Subsequently, a preliminary t-distributed stochastic neighbor embedding (t-SNE) was performed for synovial and blood samples using *RunPCA* and *RuntSNE* functions with default parameter settings. By manual inspection, 54 cells were identified as monocytic contamination based on separate clustering and detection of marker transcripts (e.g., *CD14* and *ITGAM* among others, see cluster 6 shown in Extended Fig. 2a) as well as a cluster representing FOXP3-expressing regulatory T cells (483 cells; cluster 5 shown in Extended Fig. 2a) among synovial CD4^+^CD45RO^+^CD25^−^ T cells. The respective clusters were removed from further analysis of synovial Tcon (Extended Fig. 2b). The filtered set comprised six paired PB and SF samples with 13,479 and 12,481 Tcon, respectively. In addition, Tcon from patient 3 did not have a corresponding PB sample and included 1,275 synovial Tcon (summing up to 13,756 synovial Tcon in total).

In addition, the filtered set contained 10,872 and 10,157 CD4^+^CD45RO^+^CD127^lo^CD25^+^ (Treg) cells, as well as 12,111 and 14,019 CD8^+^CD45RO^+^ T cells from the PB and the SF, respectively.

### Single-cell transcriptome profiling

Single-cell transcriptome profiling was performed using the Seurat R package (version 3.1.1). In particular, samples from the PB and the SF were filtered as described above and integrated (combined) with the functions *FindIntegrationAnchors* and *IntegrateData* as proposed by Rahul Satija and colleagues (Tim Stuart et al.) and thereby batch-corrected^101^. As input for the *FindIntegrationAnchors*, the 800 most variable features were determined within each sample. A list of unique variable features from different samples (SF Tcon, PB Tcon, SF Treg, PB Treg, SF CD8^+^CD45RO^+^ T cells and PB CD8^+^CD45RO^+^ T cells) was supplied to the integration procedure. The integrated expression matrix was z-transformed by the *ScaleData* function and used to determine the first 50 principal components (PC) using *RunPCA*. Based on significant PCs (p-value < 0.001 as determined by *JackStraw* and *ScoreJackStraw*) t-distributed stochastic neighbor embedding was performed using *RuntSNE*, with a perplexity of 50 and a theta value of 0. Finally, communities of cells (clusters) were annotated by the functions *FindNeighbors* and *FindClusters* using the Louvain algorithm and were based on the significant PCs and a resolution of 0.2^37^.

Regarding gene expression displayed by heatmaps, the heatmap color scales indicate the row-wise z-scores of expression means (determined by Seurat’s *AverageExpression*) of log-normalized (by Seurat’s *LogNormalize*) UMIs per cluster (*RNA assay* used). For unsupervised heatmaps, significantly differentially expressed genes were selected as follows: The expression level of cells from one cluster was compared to the expression level of cells from all the other clusters using the *FindAllMarkers* function. The statistical test used was Wilcoxon’s rank-sum test. Only genes (features) were included which were expressed by at least 10 % of cells in the respective cluster and which resulted in an adjusted p-value (Bonferroni correction) lower or equal to 0.05.

Cell-wise module scores for gene sets were generated by Seurat’s *AddModuleScore* function with default parameters and using the RNA assay.

Feature plots (gene expression level on t-SNE map) show log normalized (Seurat’s *LogNormalize* function) UMIs with a color scale cutoff at the 95^th^ percentile.

### Single-cell TCR repertoire profiling

The Immune-Profiling analysis by cellranger revealed 16,733 and 15,477 Tcon from six paired PB and SF samples, respectively, as well as additional 1,628 cells from the SF from a donor whose blood sample was not sequenced. For further analysis, solely cells with annotations of the v(d)j gene and cdr3 region for both TCR alpha and beta chain were considered. For cells with ambiguous alpha or beta chain annotations, contigs with the best quality, that is, productive-ness, full-length and highest number of reads and UMIs were chosen. Cells were integrated with the transcriptome analysis (including quality assessment and filtering), revealing 7,098 out of 13,479 cells and 7,403 out of 12,481 cells from six paired Tcon blood and synovial fluid samples, respectively, plus 1,042 out of 1,275 cells from synovial Tcon with no corresponding blood sample. For overlaps between PB and SF cells with no corresponding blood sample were not considered. The clonotypes were defined by the donor origin, the v(d)j genes and cdr3nt annotations for both the alpha and beta chain. Overlaps were defined by the ratio of cells in a cluster with clonotypes shared by a cluster being compared. Overlaps between two clusters were compared to overlaps between these two clusters with randomized cluster annotations. Overlaps of clusters were considered to be significant when not more than 5 out 1000 randomized overlaps were higher than the experimentally measured overlap of clusters.

### Monocle trajectories, GSEA and Shortest-Path analysis

In order to determine a pseudotime cell trajectory using the monocle-package (version 2.12) by Cole Trapnell, processed data had to be transferred from the Seurat object to monocle’s CellDataSet object. The data provided to monocle were the expression matrix of Seurat’s integrated assay. *ExpressionFamily* was set to “uninormal“, since UMI-processing has been done in Seurat before. In *setOrderingFilter*, all genes of the integrated assay were selected as *ordering_genes*. In the function *reduceDimension*, parameter “norm_method” was set to “none“, “reduction_method” set to “DDRTree”, “pseudo_expr” to “0” and “max_components” set to “6”. Other arguments were left default. Cells were then arranged by *orderCells* with default settings apart from manually setting the “root_state” to the state that mostly corresponded to Seurat’s Cluster 0.

The gene set enrichment analysis (GSEA) was performed based on the CERNO test analog to the tmod R package using pre-ranked genes and Reactome pathways as gene sets^51,102–104^. In particular, for each single cell, genes were ranked by the difference to the mean expression among all synovial Tcon. Gene sets with p-values ≤ 0.05 and a false discovery rate (FDR) ≤ 0.25 were considered as significant and projected onto the Monocle-based trajectory. Visualized is the area under the curve (AUC)-values for cells showing significant enrichment.

For the Shortest-Path analysis an undirected, weighted graph was generated for cells with common TCR annotations, based on Monocle and assuming that each cell could represent a reference intermediate state. Each cell was connected to its 10 nearest neighbors based on Euclidian distance. The Shortest-Path was computed for each pair using the Dijkstra algorithm. The importance of a cell was judged by the Betweenness-Centrality. Significant paths between cells with a Betweenness-Centrality greater than 0.1 * n * (n−1), with n being the number of cells, were visualized. The direction of an edge was judged by the higher centrality value.

### Statistical analyses

All statistical tests are specified in the figure legends of the respective analyses, as well as in the respective material and methods section. Statistical calculations were performed with GraphPad Prism version 5.04/7.04 and R version 3.6.1 if not specified otherwise.

## Acknowledgements

This work was supported by the state of Berlin and the “European Regional Development Fund” (ERDF 2014–2020, EFRE 1.8/11, Deutsches Rheuma-Forschungszentrum to M.F.M.), Deutsche Forschungsgemeinschaft through DFG priority program 1468 IMMUNOBONE and the TR130 (to A.R. and H.D.C.) and by the European Research Council through the Advanced Grant IMMEMO (ERC-2010-AdG.20100317 Grant 268978 to AR), the Innovative Medicines Initiative 2 Joint Undertaking under grant agreement No 777357, the Rheumastiftung (to H.D.C.) and the Leibniz Science Campus Chronic Inflammation (www.chronische-entzuendung.org). P.M. has been supported by EUTRAIN, a FP7 Marie Curie Initial Training Network for Early Stage Researchers funded by the European Union (FP7-PEOPLE-2011-ITN-289903). H.D.C. is funded by the Dr. Rolf M. Schwiete Foundation. M.A.M. has been supported by the Einstein Foundation Berlin (EP-2017-393). C.M.S. is supported by the Deutsche Forschungsgemeinschaft (DFG; EN 924/5-1). We thank Fahd Qadir (GitHub user Dragonmasterx87) for providing code to transfer data from Seurat to Monocle. F.W. and L.L. were supported by a VIDI grant from ZonMw (91714332).

---

**Extended Data Figure 1:**
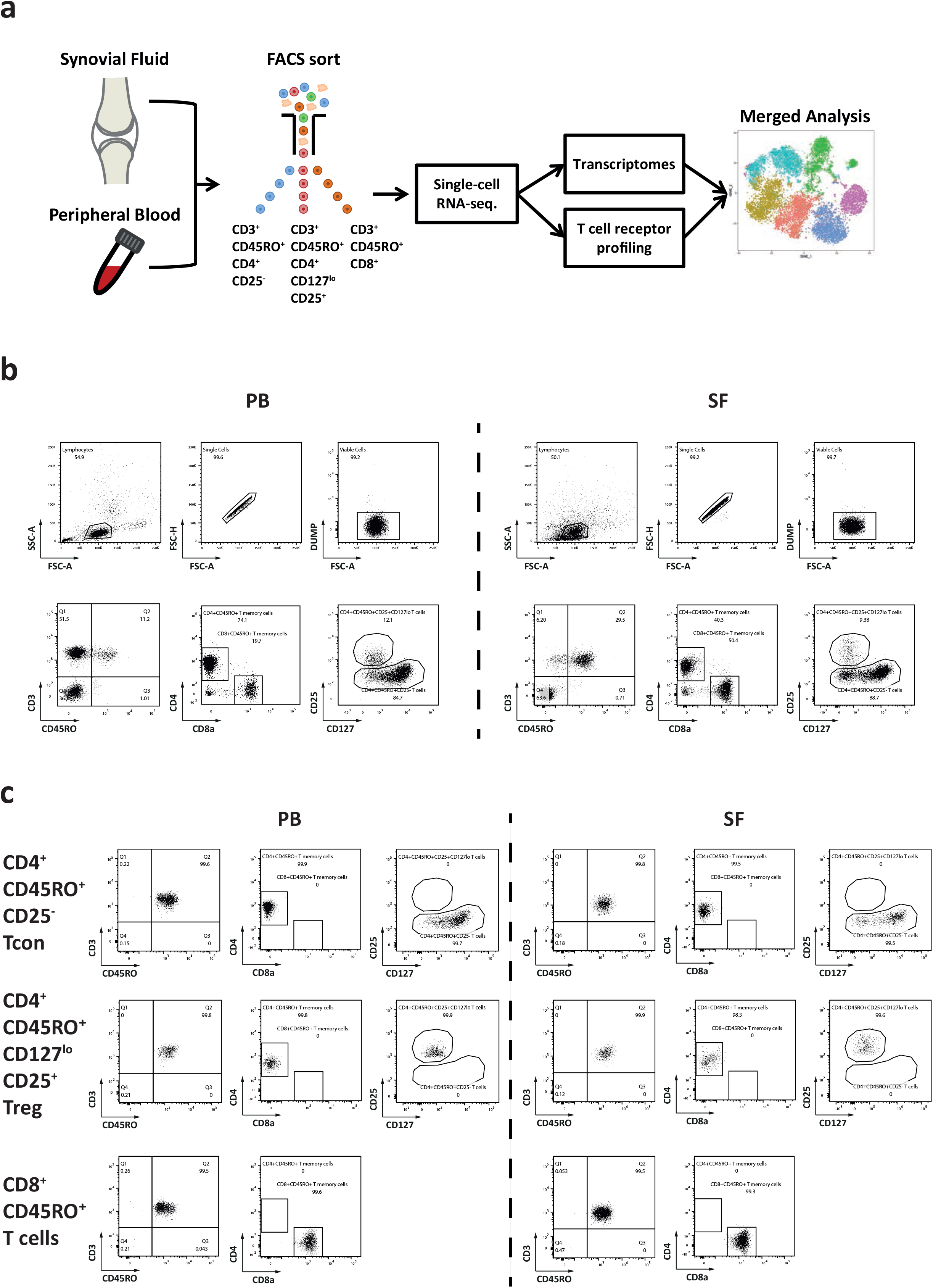
Analysis of single-cell transcriptomes and TCR repertoires of antigen-experienced (memory) T cell subsets. **a**, Schematic of the study design to analyze single cell transcriptomes and the TCR repertoire of JIA patients. CD45RO^+^ (antigen-experienced or memory) T lymphocytes were isolated from the synovial fluid (SF) and the peripheral blood (PB) to generate pairwise single-cell transcriptomic and TCR repertoire data of JIA patients. Exceptions for pairwise analysis were made for patient 3, whose blood we were not able to analyze with regards to Tcon and CD8^+^CD45RO^+^ T cells, as well as for patient 6, whose SF Treg cells we were not able to sequence. **b**, Representative dot plots showing the gating strategy and hierarchy to isolate populations of CD4^+^CD45RO^+^CD25^−^ Tcon, CD4^+^CD45RO^+^CD127^lo^CD25^+^ Treg and CD8^+^CD45RO^+^ T cells from the PB and the SF of JIA patients by flow cytometry. **c**, Representative dot plots of flow-sorted Tcon, Treg and CD8^+^CD45RO^+^ T cells. Following flow sort, purified cells were subjected to 10X Genomics-based single-cell sequencing.

**Extended Data Figure 2:**
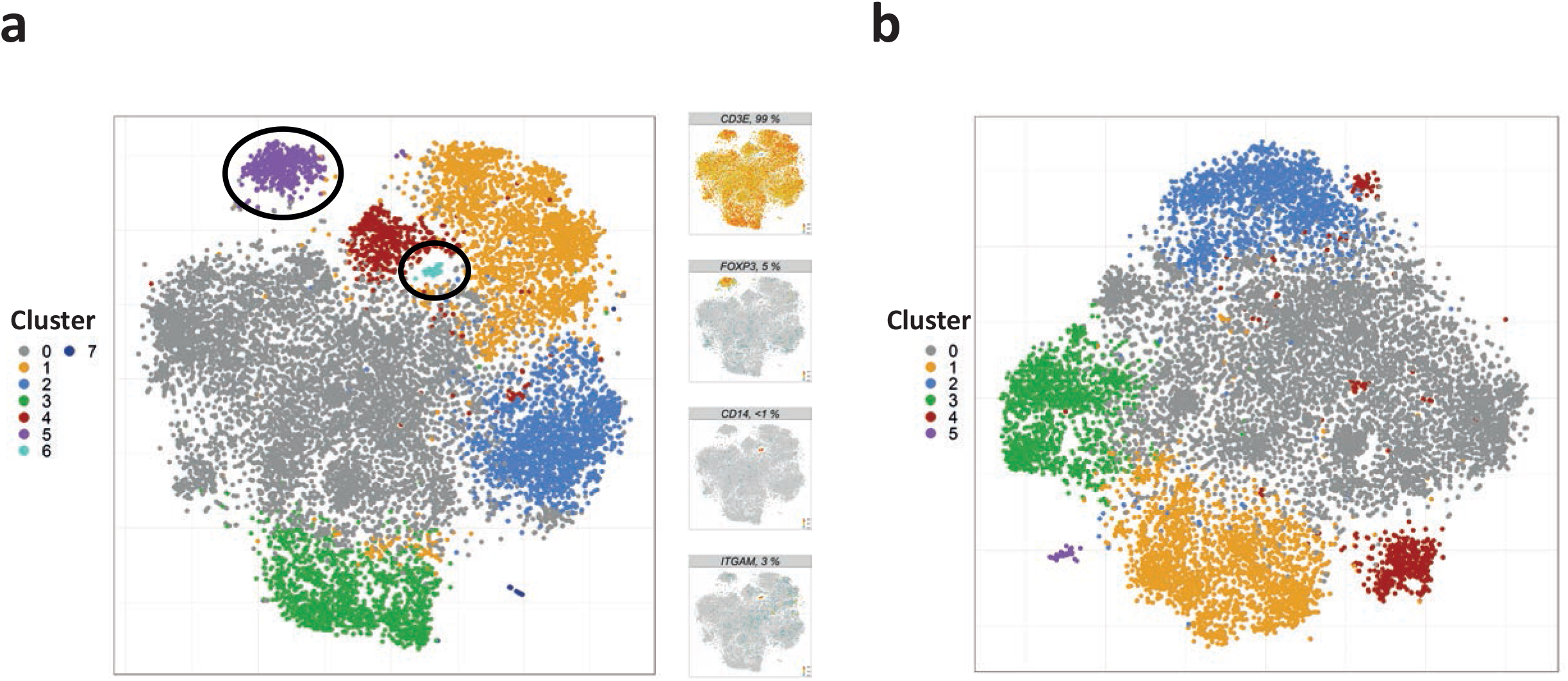
Clustering and t-SNE projection of synovial Tcon following removal of contaminating cells. **a**, Synovial CD4^+^CD45RO^+^CD25^−^ T cells were clustered and projected on a t-SNE map. **b**, A t-SNE map of synovial Tcon. The t-SNE map was generated from CD4^+^CD45RO^+^CD25^−^ T cells after removal of two clusters containing *FOXP3*^+^ regulatory T cells and *CD14*^+^ monocytes.

**Extended Data Figure 3:**
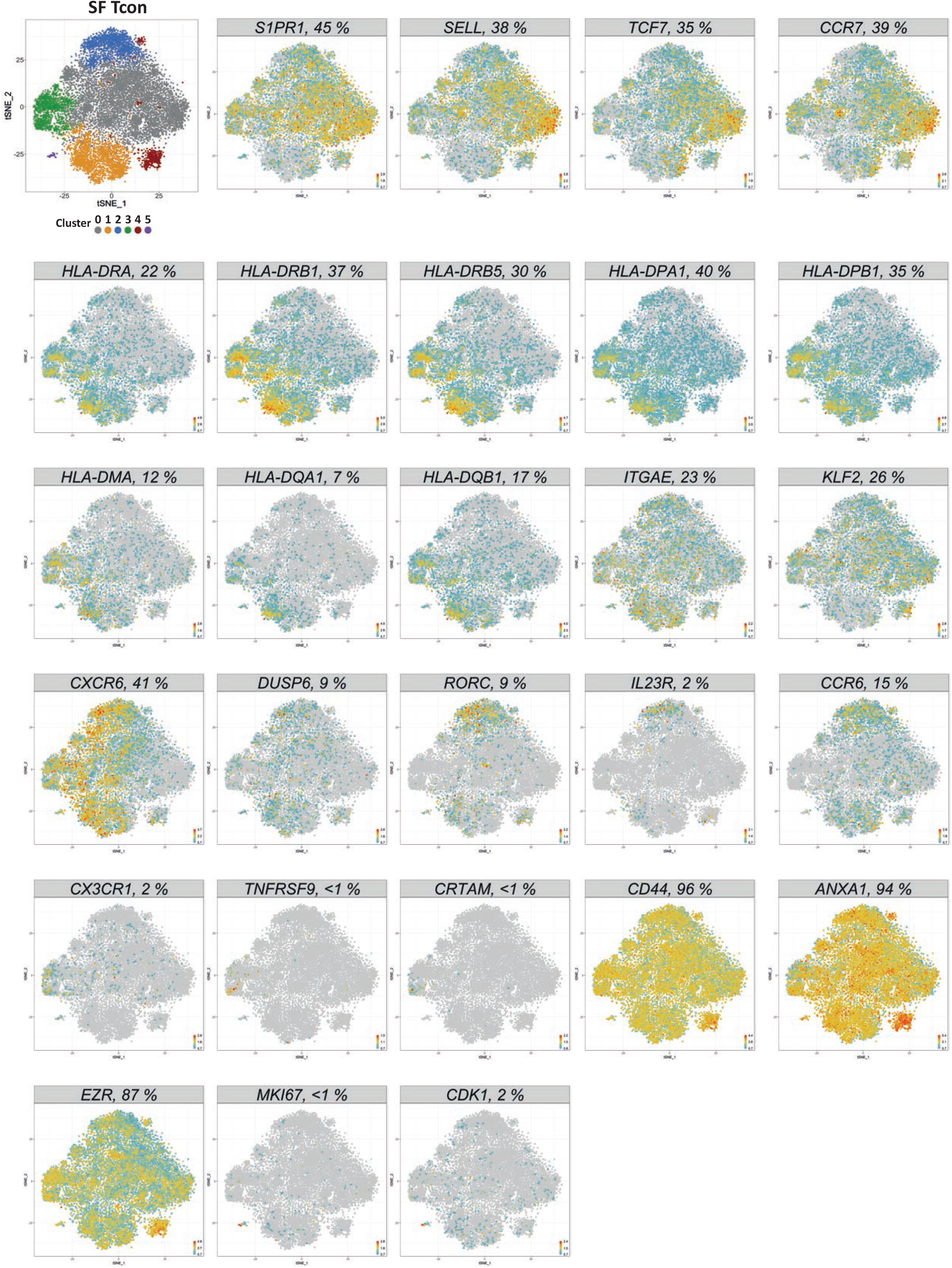
Genes that are expressed by different subsets of Tcon from inflamed joints of JIA patients. A t-SNE map and feature plots with indicated genes expressed by different subsets of synovial Tcon that were isolated from inflamed joints of JIA patients. Percentages in the header bars of the feature plots display the frequencies of Tcon that express the indicated genes.

**Extended Data Figure 4:**
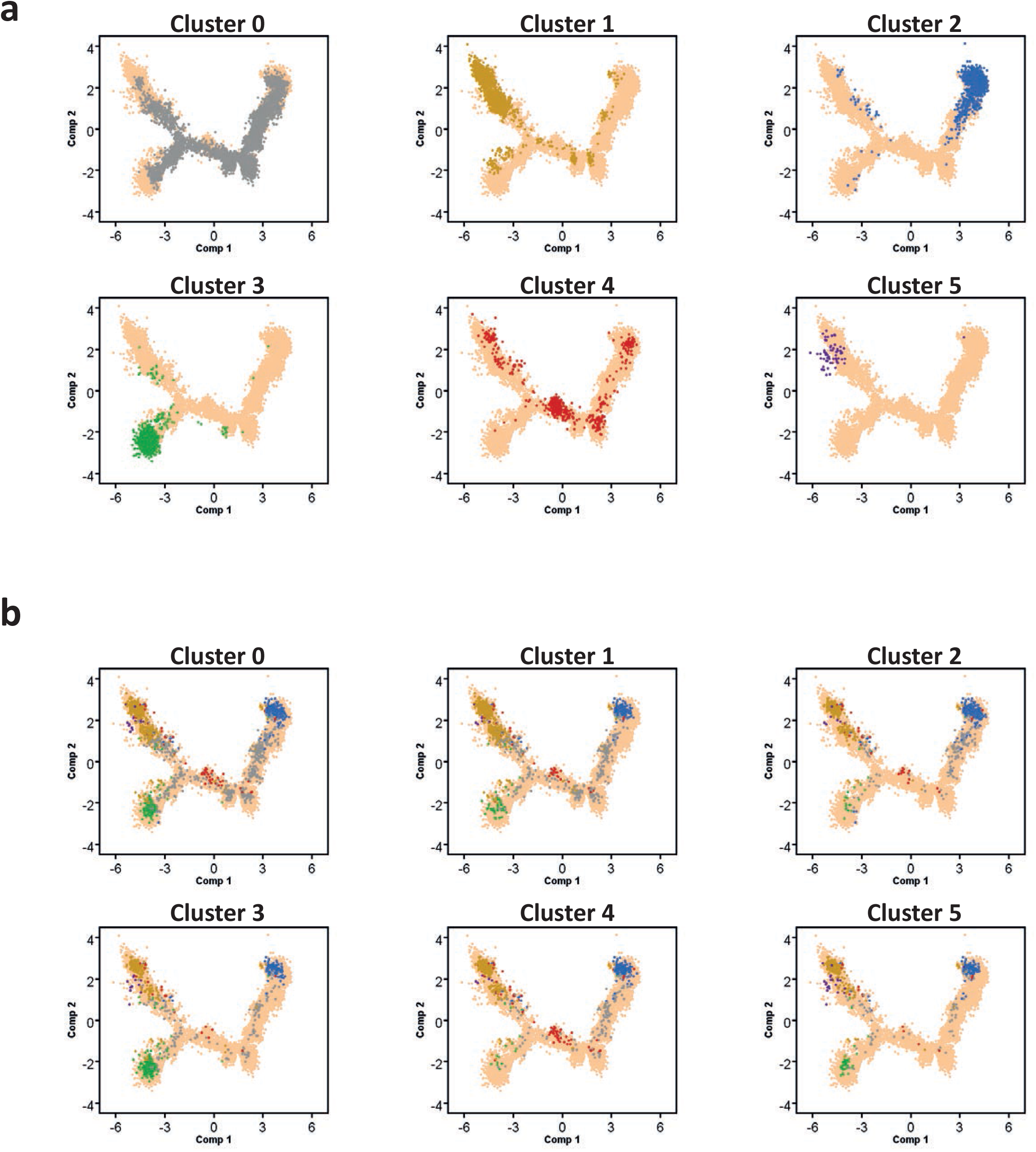
Distribution of cells and expanded clones among synovial Tcon clusters. Trajectories of synovial Tcon were generated by Monocle as described in **Fig. 2**. **a**, Depicted is the distribution of synovial Tcon among the trajectory according to their cluster association. **b**, Shown are expanded clones that were found in the indicated clusters and their distribution among the other clusters of synovial Tcon. In **a** and **b**, Each cell is represented as a dot and colored according to its Tcon cluster color: i.e. grey (cluster 0), orange (cluster 1), blue (cluster 2), green (cluster 3), red (cluster 4), violet (cluster 5). Cells that are not highlighted are depicted as yellow in the background to outline the shape of the trajectory.

**Extended Data Figure 5:**
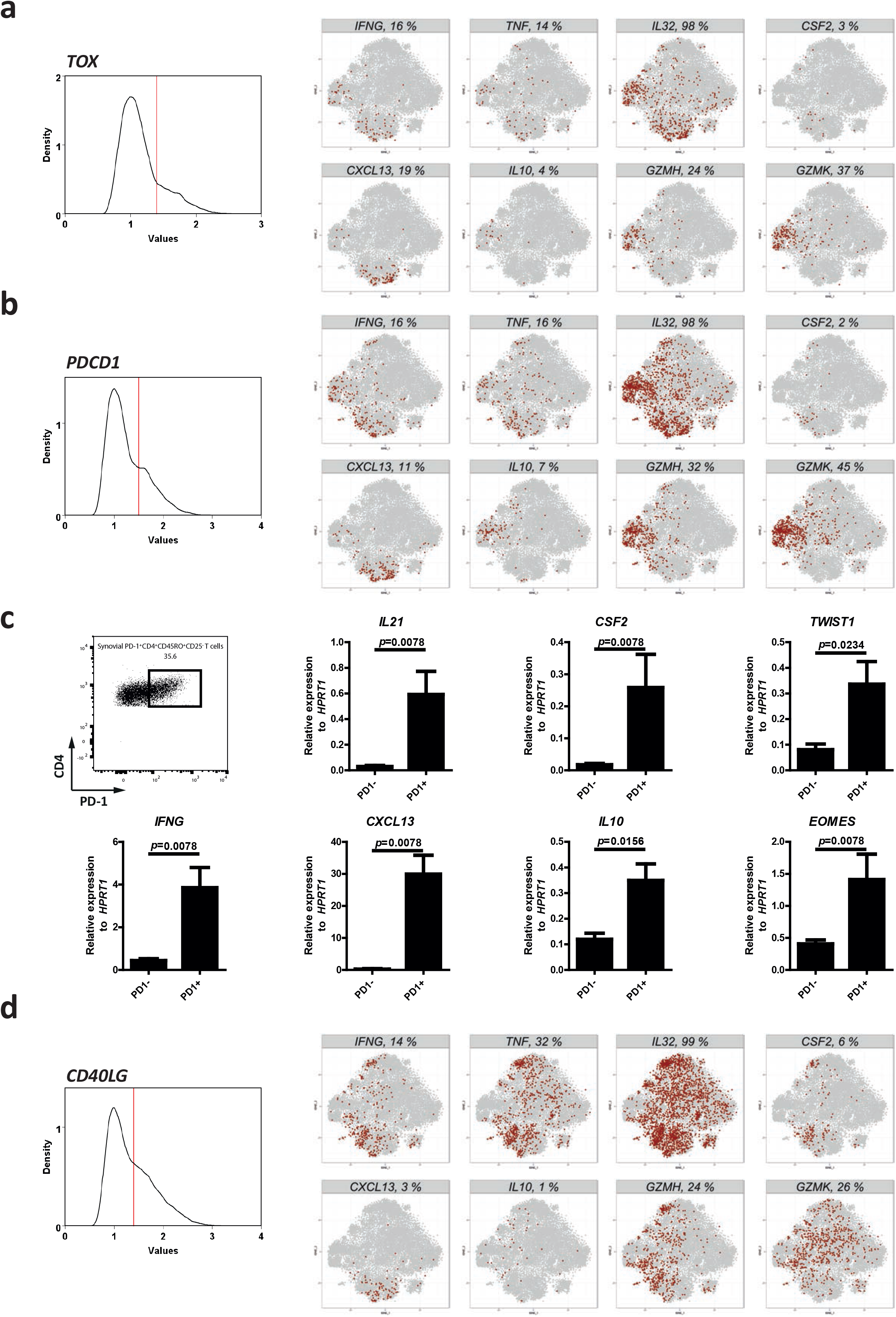
Synovial PD-1^+^*TOX*^+^ Tcon of JIA patients co-express genes of activation. **a**, **b** and **d**, Displayed are feature plots of synovial Tcon that were “gated” (histograms) on *TOX*^+^ (cells ≥ 1.4 normalized UMI counts) (**a**), *PDCD1*^+^ (cells ≥ 1.4 normalized UMI counts) (**b**), and *CD40LG*^+^ (cells ≥ 1.5 normalized UMI counts) (**d**) synovial Tcon of JIA patients (n=7). Feature plots of the gated cells indicate co-expression of the indicated functional genes (present in red, or absent in grey). The header bars of the feature plots indicate the percentages of cells expressing the indicated genes among *TOX*^+^ (**a**), *PDCD1*^+^ (**b**) and among *CD40LG*^+^ (**d**) synovial Tcon. **c**, PD-1^+^ and PD-1^−^ Tcon of JIA patients were sorted by FACS (n=8, among these, 5 patients were diagnosed with oligoarticular JIA, 2 patients were diagnosed with psoriatic arthritis and 1 patient was diagnosed with polyarticular JIA). Afterwards, the *in situ* expression of the indicated functional genes including cytokine genes, chemokine genes and genes encoding the transcription factors *TWIST1* and *EOMES* by the sorted cells was measured by qPCR. Statistical significance was tested by a two-tailed Mann Whitney test.

**Extended Data Figure 6:**
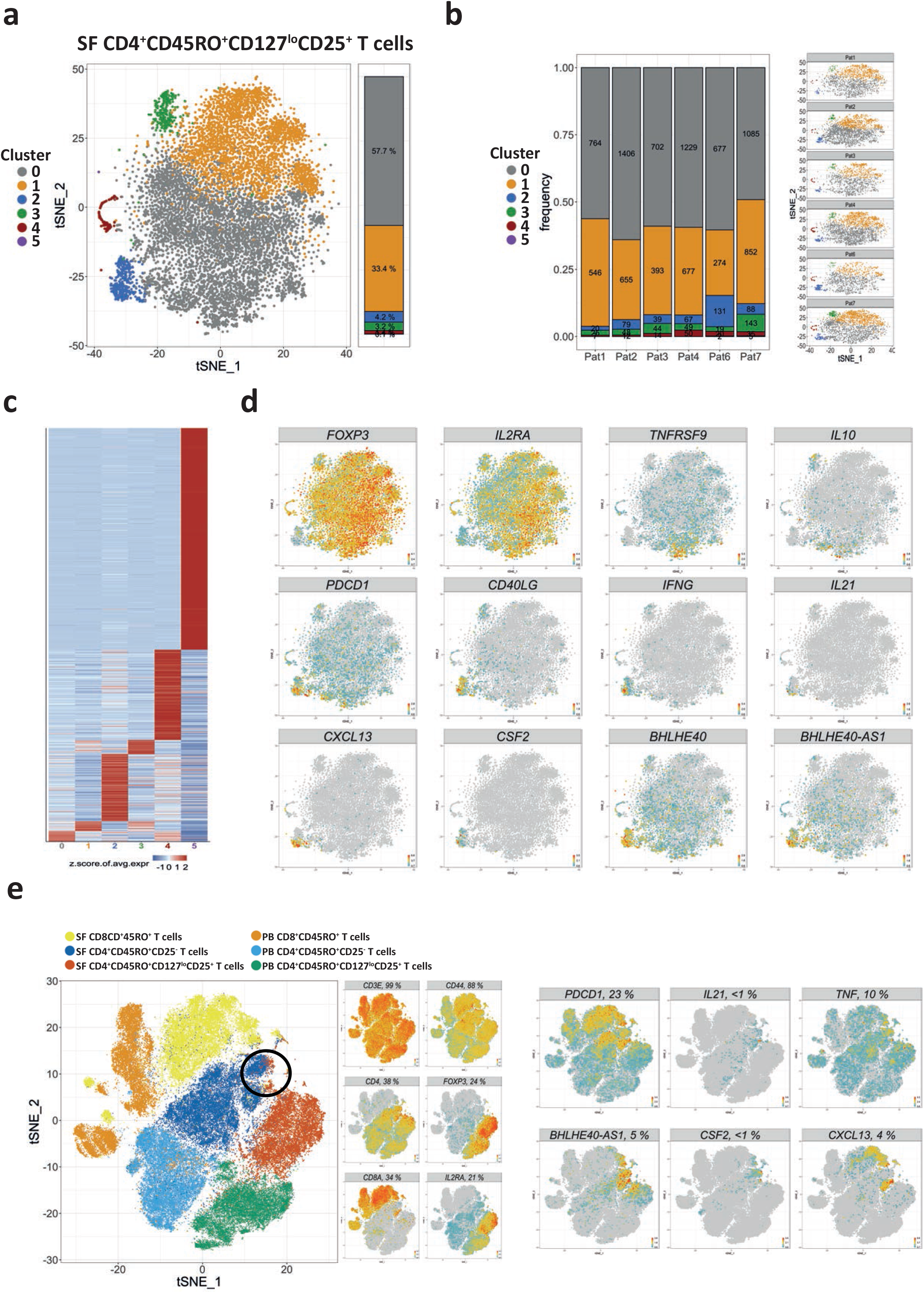
Synovial CD4^+^CD45RO^+^CD127^lo^CD25^+^ T cells contain cells that resemble pro-inflammatory cells of synovial Tcon cluster 1. To assess the heterogeneity of Treg, 10,157 CD4^+^CD45RO^+^CD127^lo^CD25^+^ T cells were purified from inflamed joints of 6 JIA patients by FACS and clustered following single-cell sequencing. **a**, A combined t-SNE map depicting the clusters of CD4^+^CD45RO^+^CD127^lo^CD25^+^ T cells from the inflamed joints of 6 JIA patients. **b**, Quantification and t-SNE projection of the cluster distribution among synovial CD4^+^CD45RO^+^CD127^lo^CD25^+^ T cells of the 6 different patients. **c**, Heatmap showing the z-score normalized mean expression of 838 cluster-specific genes. **d**, Feature plots of indicated genes showing that cluster 2 harbors pro-inflammatory T cells resembling cells of CD4^+^ Tcon cluster 1. **e**, A t-SNE map depicting all T memory cells from the PB and the SF, as well as feature plots of indicated genes. The area on the t-SNE map where synovial cells of Treg cluster 5 are located adjacent to synovial cells of Tcon cluster 1 is highlighted with a black circle. Heatmap color scales indicate row-wise z-scores of cluster-means of log-normalized expression values. Signature genes were tested for significant differential expression by cells of a cluster in comparison to cells from all other clusters by Wilcoxon’s rank sum test with Bonferroni correction. Cutoff was a minimum of 5 % expressing cells in the respective cluster and an adjusted p-value ≤ 0.05.

**Extended Data Figure 7:**
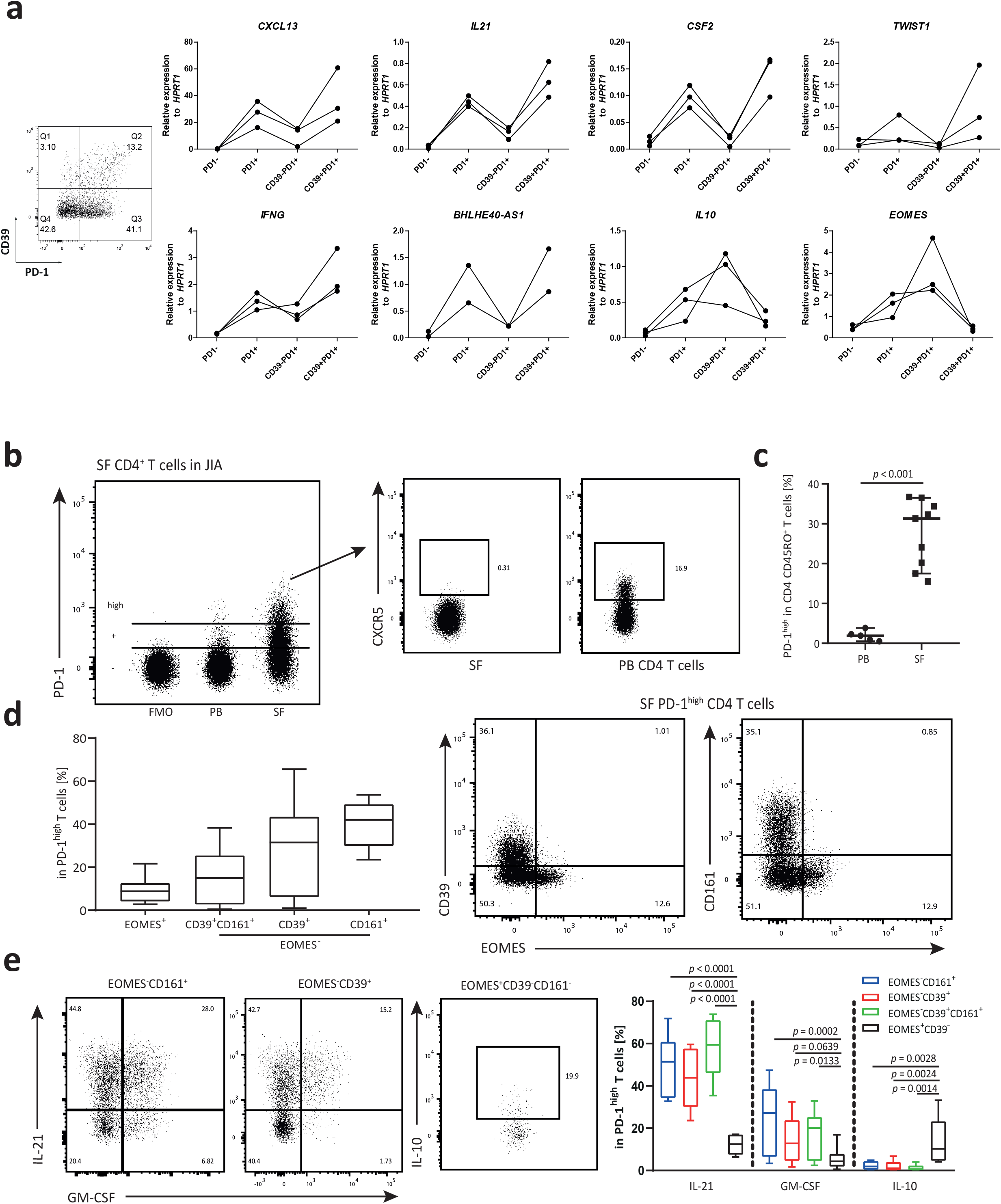
Presence of distinct PD1^high^ T cell subsets that drive or regulate chronic inflammation in JIA. **a**, Tcon from the SF of JIA patients were sorted according to PD1 and CD39 expression. Then the sorted cells were lysed to isolate the RNA and to determine *in situ* expression of indicated genes by qPCR. Shown is a representative FACS plot of PD-1 and CD39 expression among Tcon, as well as plots that quantify the gene expression of depicted genes (n=3, these patients were also included among the patients shown in Extended Data Fig. 5; 2 of the 3 patients were diagnosed with oligoarticular JIA and 1 patient was diagnosed with psoriatic arthritis. Results for BHLHE40-AS1 contain data from n=2 patients). **b-e**, SF T cells of JIA patients and T cells from the PB of both JIA patients and healthy controls were stimulated and stained for analysis by flow cytometry. **b**, Representative FACS plots of PD-1-stained CD4^+^ T cells from the SF and the PB including a fluorescent minus one (FMO) control. Subsequent gating shows absence of CXCR5 expression among PD-1^high^CD4^+^ T cells in the SF. **c**, Frequencies of PD-1^high^ T cells within CD4^+^CD45RO^+^ T cells of the PB (n=5) and the SF (n=9). **d**, Frequencies of EOMES^+^, EOMES^−^CD39^+^, EOMES^−^CD161^+^, and EOMES^−^CD39^+^CD161^+^ cells within synovial PD-1^high^ CD4^+^ T cells of JIA patients (n=9). Representative FACS plots showing mutually exclusive expression of CD39 and EOMES, as well as mutually exclusive expression of CD161 and EOMES among synovial PD-1^high^CD4^+^ T lymphocytes of JIA patients. **e**, Representative FACS plots showing the expression of IL-21 and GM-CSF by EOMES^−^CD161^+^ and EOMES^−^CD39^+^ cells, as well as the expression of IL-10 by EOMES^+^CD39^−^CD161^−^ cells among SF CD4^+^PD1^high^ T cells. Boxplots show the frequencies of IL-21-, GM-CSF- and IL-10-expressing cells within EOMES^−^CD161^+^ (blue), EOMES^−^CD39^+^ (red), EOMES^−^CD39^+^CD161^+^ (green) and EOMES^+^CD39^−^CD161^−^ (black) cells among synovial PD-1^high^CD4^+^ T cells of JIA patients (n=6-7). Data in **c**, **d** and **e** are representative of 2 independent experiments. Statistical significance in **c** was determined with an unpaired Student’s t test. Statistical significance in **e** was determined with a one-way ANOVA with equal variance assumption and a Dunnett post-hoc test; the boxplots represent the median with 10-90 percentiles.

**Extended Data Figure 8:**
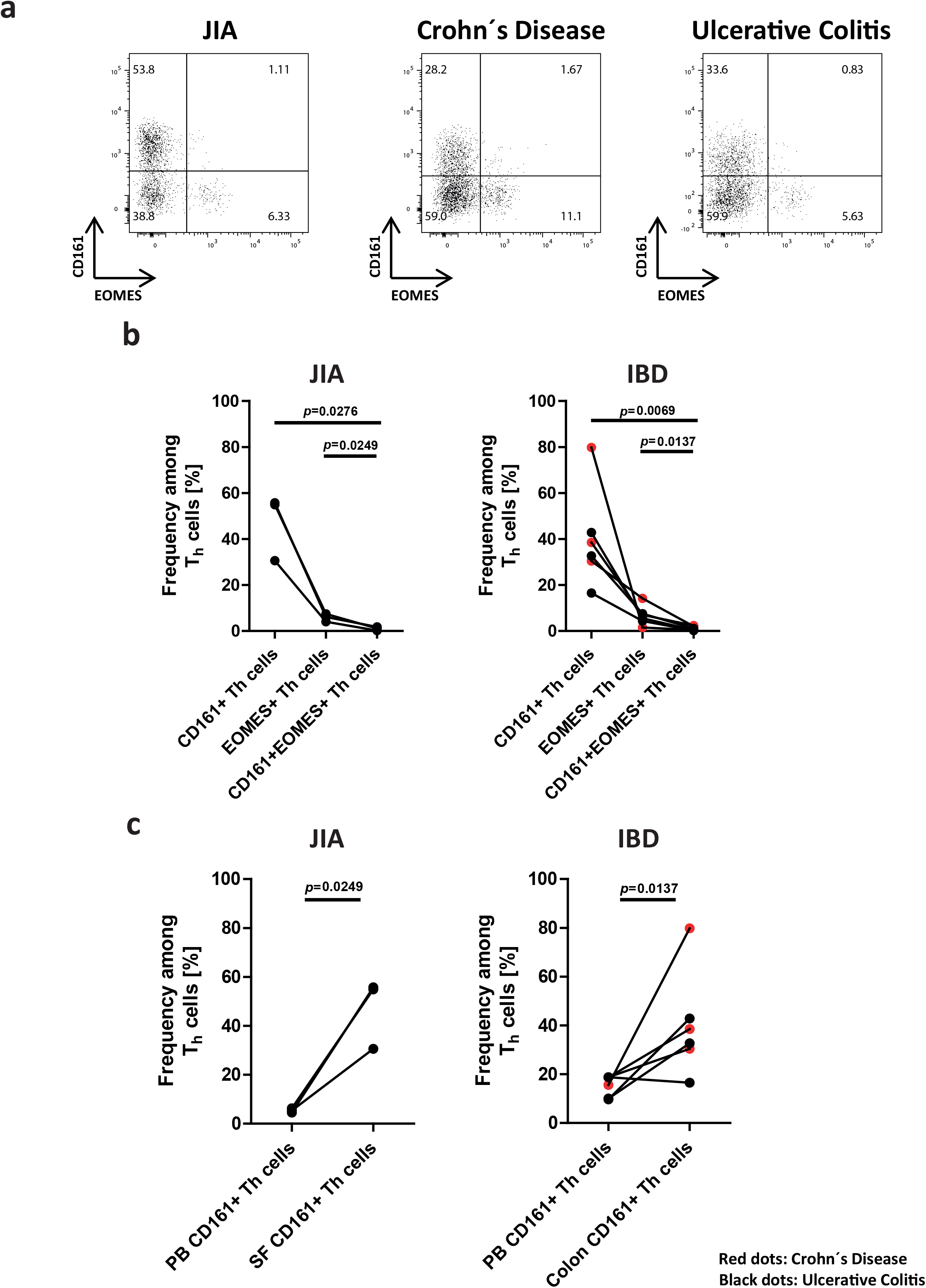
Distinct T_h_ cell subsets at inflamed sites of multiple chronic inflammatory diseases can be distinguished by expression of CD161 and EOMES. T_h_ cells were isolated from the SF and the PB of JIA patients (n=3). In addition, T_h_ cells were purified from biopsies of inflamed colons from patients with inflammatory bowel disease (IBD), i.e. with Crohn’s diseases (n=3) and ulcerative colitis (n=3). Purified T_h_ cells were stimulated and analyzed by flow cytometry. **a**, Representative FACS plots showing T_h_ cell subsets expressing either CD161 or EOMES. **b**, Frequencies of T_h_ cells from inflamed joints of JIA patients (n=3) or from inflamed colons of IBD patients (n=6) that express CD161, EOMES or both CD161 and EOMES. Red dots in the graph of IBD patients quantify T_h_ cell frequencies of patients with Crohn’s disease (n=3), while black dots represent samples from patients with ulcerative colitis (n=3). Statistical significance was determined by a paired two-tailed Student’s t test. **c**, Frequencies of CD161^+^ cells among T cells from the PB or the SF of JIA patients (n=3) and among T_h_ cells isolated from the PB or from colon biopsies of IBD patients (n=6). Red dots in the graph of IBD patients quantify cell frequencies of 3 Crohn’s disease patients, while black dots represent samples from 3 patients with ulcerative colitis. Statistical significance was determined by a paired two-tailed Student’s t test.

**Extended Data Figure 9:**
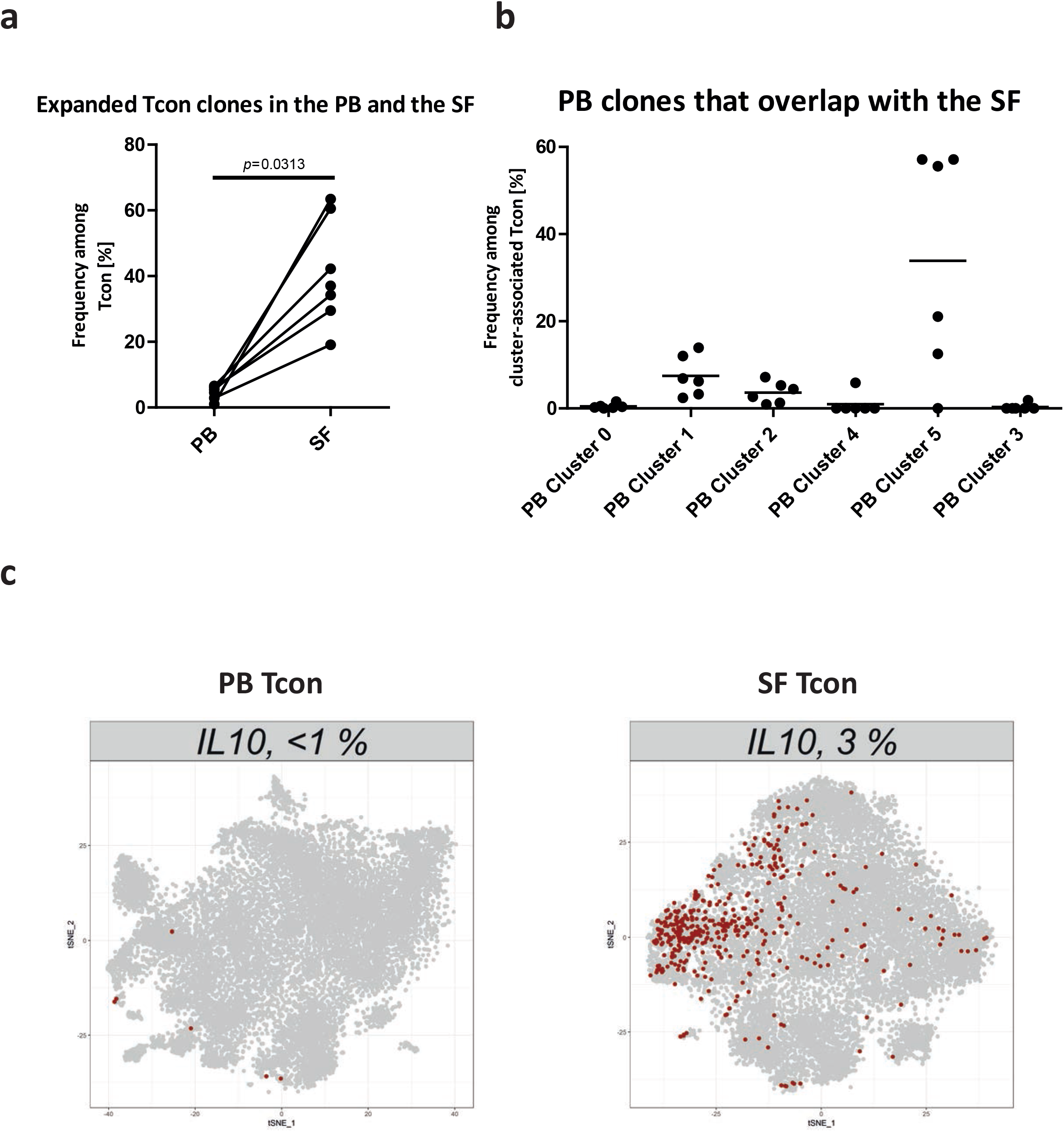
Clonal overlap and *IL10* expression of CD4^+^ PB Tcon. **a**, The graph shows the frequencies of expanded Tcon clones from 6 paired SF and PB samples and an additional SF patient without corresponding PB sample. **b**, Frequencies of expanded T cell clones among all PB Tcon clusters are shown. **c**, Feature plots showing *IL10*-expressing Tcons from the PB and the SF. Red dots represent Tcon that express *IL10*, while grey dots represent Tcon for which no *IL10* transcripts were detected. Statistical significance in a was tested with a Wilcoxon matched-pairs signed rank test.

**Extended Data Figure 10:**
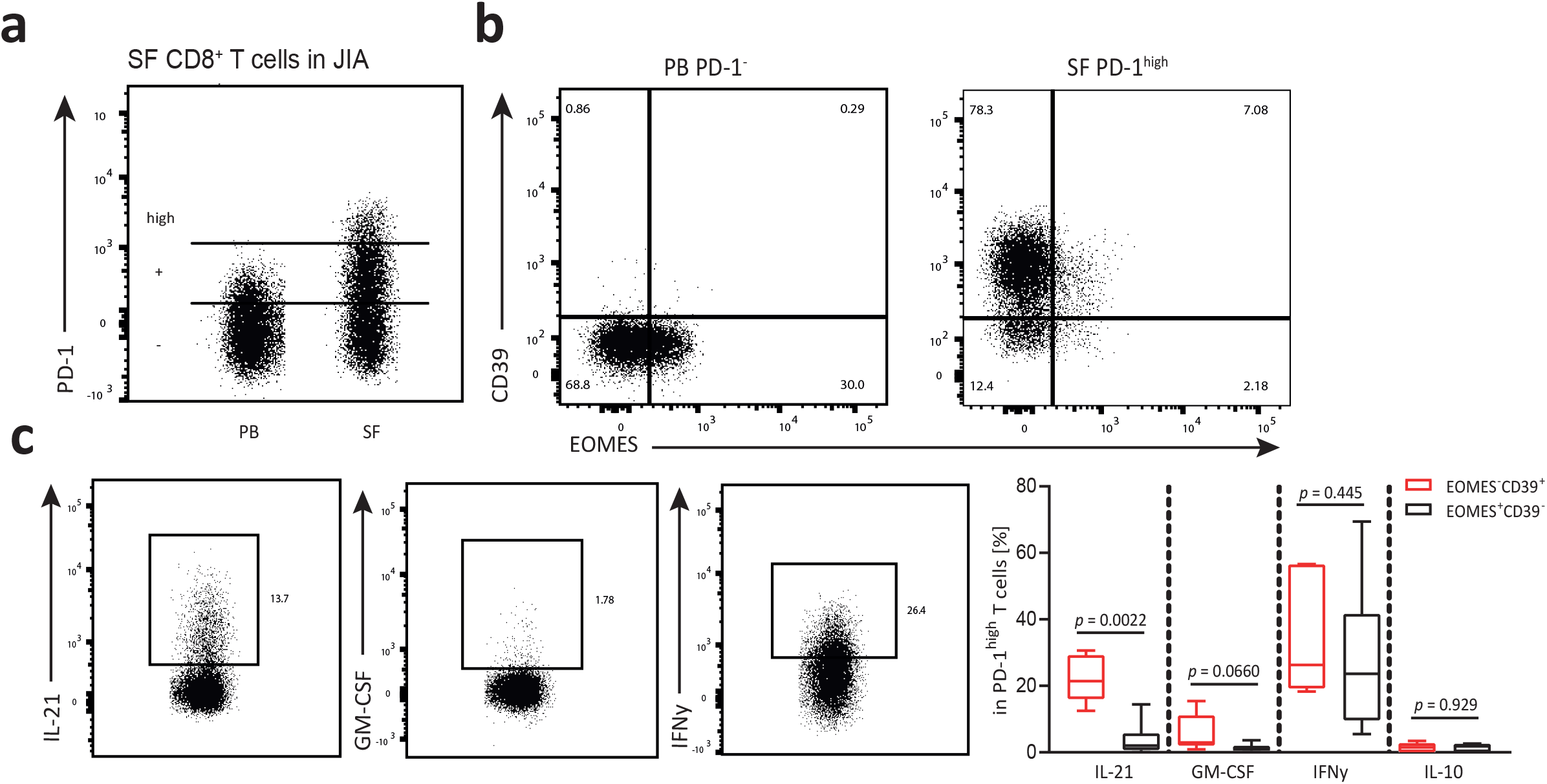
Identification of a T_h_ cell-like CD8^+^ T cell population that is present in inflamed joints of JIA patients. **a**, Representative FACS plots showing PD-1 expression among SF and PB CD8^+^FOXP3^−^ T cells. **b**, Representative FACS plots displaying the gating of CD39^+^ and EOMES^+^ showing a dichotomy for both PB PD-1^−^ and SF PD-1^high^ CD8^+^FOXP3^−^ T cells. **c**, Representative FACS plots showing the expression of IL-21, GM-CSF and IFN-γ by SF CD8^+^FOXP3^−^PD1^high^EOMES^−^CD39^+^ T cells. Frequencies of IL-21, GM-CSF, IFN-γ and IL-10 within the EOMES^−^CD39^+^ (red) and EOMES^+^CD39^−^ (black) cells within SF CD8^+^FOXP3^−^ PD1^high^ T cells (n=6-7). Data in **c** are representative of 2 independent experiments. Statistical significance in **c** was determined with a paired Student’s t-test.

## References

1. Van Boxel, J. A. & Paget, S. A. Predominantly T-cell infiltrate in rheumatoid synovial membranes. N Engl J Med 293, 517–520, doi:10.1056/NEJM197509112931101 (1975).

2. Duke, O., Panayi, G. S., Janossy, G. & Poulter, L. W. An immunohistological analysis of lymphocyte subpopulations and their microenvironment in the synovial membranes of patients with rheumatoid arthritis using monoclonal antibodies. Clin Exp Immunol 49, 22–30 (1982).

3. Shale, M., Schiering, C. & Powrie, F. CD4(+) T-cell subsets in intestinal inflammation. Immunol Rev 252, 164–182, doi:10.1111/imr.12039 (2013).

4. Abraham, C. & Cho, J. H. Inflammatory bowel disease. N Engl J Med 361, 2066–2078, doi:10.1056/NEJMra0804647 (2009).

5. Kaser, A., Zeissig, S. & Blumberg, R. S. Inflammatory bowel disease. Annu Rev Immunol 28, 573–621, doi:10.1146/annurev-immunol-030409-101225 (2010).

6. Wedderburn, L. R., Robinson, N., Patel, A., Varsani, H. & Woo, P. Selective recruitment of polarized T cells expressing CCR5 and CXCR3 to the inflamed joints of children with juvenile idiopathic arthritis. Arthritis Rheum 43, 765–774, doi:10.1002/1529-0131(200004)43:4<765::AID-ANR7>3.0.CO;2-B (2000).

7. Imam, T., Park, S., Kaplan, M. H. & Olson, M. R. Effector T Helper Cell Subsets in Inflammatory Bowel Diseases. Front Immunol 9, 1212, doi:10.3389/fimmu.2018.01212 (2018).

8. McInnes, I. B. & Schett, G. The pathogenesis of rheumatoid arthritis. N Engl J Med 365, 2205–2219, doi:10.1056/NEJMra1004965 (2011).

9. Chemin, K., Gerstner, C. & Malmstrom, V. Effector Functions of CD4+ T Cells at the Site of Local Autoimmune Inflammation-Lessons From Rheumatoid Arthritis. Front Immunol 10, 353, doi:10.3389/fimmu.2019.00353 (2019).

10. Spreafico, R. et al. Epipolymorphisms associated with the clinical outcome of autoimmune arthritis affect CD4+ T cell activation pathways. Proc Natl Acad Sci U S A 113, 13845–13850, doi:10.1073/pnas.1524056113 (2016).

11. Emmrich, J., Seyfarth, M., Fleig, W. E. & Emmrich, F. Treatment of inflammatory bowel disease with anti-CD4 monoclonal antibody. Lancet 338, 570–571, doi:10.1016/0140-6736(91)91133-f (1991).

12. Horneff, G., Burmester, G. R., Emmrich, F. & Kalden, J. R. Treatment of rheumatoid arthritis with an anti-CD4 monoclonal antibody. Arthritis Rheum 34, 129–140, doi:10.1002/art.1780340202 (1991).

13. Stronkhorst, A. et al. CD4 antibody treatment in patients with active Crohn’s disease: a phase 1 dose finding study. Gut 40, 320–327, doi:10.1136/gut.40.3.320 (1997).

14. Goronzy, J. J. & Weyand, C. M. Long-term immunomodulatory effects of T lymphocyte depletion in patients with systemic sclerosis. Arthritis Rheum 33, 511–519, doi:10.1002/art.1780330408 (1990).

15. Chang, H. D. & Radbruch, A. Targeting pathogenic T helper cell memory. Ann Rheum Dis 70 Suppl 1, i85–87, doi:10.1136/ard.2010.140954 (2011).

16. Wu, C. et al. Induction of pathogenic TH17 cells by inducible salt-sensing kinase SGK1. Nature 496, 513–517, doi:10.1038/nature11984 (2013).

17. Rao, D. A. et al. Pathologically expanded peripheral T helper cell subset drives B cells in rheumatoid arthritis. Nature 542, 110–114, doi:10.1038/nature20810 (2017).

18. Galli, E. et al. GM-CSF and CXCR4 define a T helper cell signature in multiple sclerosis. Nat Med 25, 1290–1300, doi:10.1038/s41591-019-0521-4 (2019).

19. Burmester, G. R. et al. Mavrilimumab, a human monoclonal antibody targeting GM-CSF receptor-alpha, in subjects with rheumatoid arthritis: a randomised, double-blind, placebo-controlled, phase I, first-in-human study. Ann Rheum Dis 70, 1542–1549, doi:10.1136/ard.2010.146225 (2011).

20. Burmester, G. R. et al. Efficacy and safety of mavrilimumab in subjects with rheumatoid arthritis. Ann Rheum Dis 72, 1445–1452, doi:10.1136/annrheumdis-2012-202450 (2013).

21. Cook, A. D. & Hamilton, J. A. Investigational therapies targeting the granulocyte macrophage colony-stimulating factor receptor-alpha in rheumatoid arthritis: focus on mavrilimumab. Ther Adv Musculoskelet Dis 10, 29–38, doi:10.1177/1759720X17752036 (2018).

22. Becher, B., Tugues, S. & Greter, M. GM-CSF: From Growth Factor to Central Mediator of Tissue Inflammation. Immunity 45, 963–973, doi:10.1016/j.immuni.2016.10.026 (2016).

23. Niesner, U. et al. Autoregulation of Th1-mediated inflammation by twist1. J Exp Med 205, 1889–1901, doi:10.1084/jem.20072468 (2008).

24. Albrecht, I. et al. Persistence of effector memory Th1 cells is regulated by Hopx. Eur J Immunol 40, 2993–3006, doi:10.1002/eji.201040936 (2010).

25. Haftmann, C. et al. miR-148a is upregulated by Twist1 and T-bet and promotes Th1-cell survival by regulating the proapoptotic gene Bim. Eur J Immunol 45, 1192–1205, doi:10.1002/eji.201444633 (2015).

26. Maschmeyer, P. et al. Selective targeting of pro-inflammatory Th1 cells by microRNA-148a-specific antagomirs in vivo. J Autoimmun 89, 41–52, doi:10.1016/j.jaut.2017.11.005 (2018).

27. Petrelli, A. & van Wijk, F. CD8(+) T cells in human autoimmune arthritis: the unusual suspects. Nat Rev Rheumatol 12, 421–428, doi:nrrheum.2016.74 [pii] 10.1038/nrrheum.2016.74 (2016).

28. Petrelli, A. et al. Self-Sustained Resistance to Suppression of CD8+ Teff Cells at the Site of Autoimmune Inflammation Can Be Reversed by Tumor Necrosis Factor and Interferon-gamma Blockade. Arthritis Rheumatol 68, 229–236, doi:10.1002/art.39418 (2016).

29. Petrelli, A. et al. PD-1+CD8+ T cells are clonally expanding effectors in human chronic inflammation. J Clin Invest 128, 4669–4681, doi:10.1172/JCI96107 (2018).

30. Prakken, B., Albani, S. & Martini, A. Juvenile idiopathic arthritis. Lancet 377, 2138–2149, doi:S0140-6736(11)60244-4 [pii] 10.1016/S0140-6736(11)60244-4 (2011).

31. Prahalad, S. & Glass, D. N. A comprehensive review of the genetics of juvenile idiopathic arthritis. Pediatr Rheumatol Online J 6, 11, doi:1546-0096-6-11 [pii] 10.1186/1546-0096-6-11 (2008).

32. Smolen, J. S., Aletaha, D. & McInnes, I. B. Rheumatoid arthritis. Lancet 388, 2023–2038, doi:S0140-6736(16)30173-8 [pii] 10.1016/S0140-6736(16)30173-8 (2016).

33. Stokkers, P. C., Reitsma, P. H., Tytgat, G. N. & van Deventer, S. J. HLA-DR and -DQ phenotypes in inflammatory bowel disease: a meta-analysis. Gut 45, 395–401, doi:10.1136/gut.45.3.395 (1999).

34. Westendorf, K. et al. Unbiased transcriptomes of resting human CD4(+) CD45RO(+) T lymphocytes. Eur J Immunol 44, 1866–1869, doi:10.1002/eji.201344323 (2014).

35. Cossarizza, A. et al. Guidelines for the use of flow cytometry and cell sorting in immunological studies. Eur J Immunol 47, 1584–1797, doi:10.1002/eji.201646632 (2017).

36. Cossarizza, A. et al. Guidelines for the use of flow cytometry and cell sorting in immunological studies (second edition). Eur J Immunol 49, 1457–1973, doi:10.1002/eji.201970107 (2019).

37. Andrews, T. S. & Hemberg, M. Identifying cell populations with scRNASeq. Mol Aspects Med 59, 114–122, doi:10.1016/j.mam.2017.07.002 (2018).

38. Tirosh, I. et al. Dissecting the multicellular ecosystem of metastatic melanoma by single-cell RNA-seq. Science 352, 189–196, doi:10.1126/science.aad0501 (2016).

39. Nakashima, D. et al. Impaired initiation of contact hypersensitivity by FTY720. J Invest Dermatol 128, 2833–2841, doi:10.1038/jid.2008.174 (2008).

40. Cyster, J. G. & Schwab, S. R. Sphingosine-1-phosphate and lymphocyte egress from lymphoid organs. Annu Rev Immunol 30, 69–94, doi:10.1146/annurev-immunol-020711-075011 (2012).

41. Sallusto, F., Lenig, D., Forster, R., Lipp, M. & Lanzavecchia, A. Two subsets of memory T lymphocytes with distinct homing potentials and effector functions. Nature 401, 708–712, doi:10.1038/44385 (1999).

42. Nish, S. A. et al. CD4+ T cell effector commitment coupled to self-renewal by asymmetric cell divisions. J Exp Med 214, 39–47, doi:10.1084/jem.20161046 (2017).

43. Takada, K. et al. Kruppel-like factor 2 is required for trafficking but not quiescence in postactivated T cells. J Immunol 186, 775–783, doi:10.4049/jimmunol.1000094 (2011).

44. Weber, J. P. et al. ICOS maintains the T follicular helper cell phenotype by down-regulating Kruppel-like factor 2. J Exp Med 212, 217–233, doi:10.1084/jem.20141432 (2015).

45. Abbas, A. R., Wolslegel, K., Seshasayee, D., Modrusan, Z. & Clark, H. F. Deconvolution of blood microarray data identifies cellular activation patterns in systemic lupus erythematosus. PLoS One 4, e6098, doi:10.1371/journal.pone.0006098 (2009).

46. Ferenczi, K., Burack, L., Pope, M., Krueger, J. G. & Austin, L. M. CD69, HLA-DR and the IL-2R identify persistently activated T cells in psoriasis vulgaris lesional skin: blood and skin comparisons by flow cytometry. J Autoimmun 14, 63–78, doi:10.1006/jaut.1999.0343 (2000).

47. Frentsch, M. et al. Direct access to CD4+ T cells specific for defined antigens according to CD154 expression. Nat Med 11, 1118–1124, doi:10.1038/nm1292 (2005).

48. Bacher, P. et al. Regulatory T Cell Specificity Directs Tolerance versus Allergy against Aeroantigens in Humans. Cell 167, 1067–1078 e1016, doi:10.1016/j.cell.2016.09.050 (2016).

49. Ko, H. S., Fu, S. M., Winchester, R. J., Yu, D. T. & Kunkel, H. G. Ia determinants on stimulated human T lymphocytes. Occurrence on mitogen-and antigen-activated T cells. J Exp Med 150, 246–255, doi:10.1084/jem.150.2.246 (1979).

50. Holling, T. M., Schooten, E. & van Den Elsen, P. J. Function and regulation of MHC class II molecules in T-lymphocytes: of mice and men. Hum Immunol 65, 282–290, doi:10.1016/j.humimm.2004.01.005 (2004).

51. Fabregat, A. et al. The Reactome Pathway Knowledgebase. Nucleic Acids Res 46, D649–D655, doi:10.1093/nar/gkx1132 (2018).

52. Siracusa, F. et al. CD69(+) memory T lymphocytes of the bone marrow and spleen express the signature transcripts of tissue-resident memory T lymphocytes. Eur J Immunol 49, 966–968, doi:10.1002/eji.201847982 (2019).

53. Kumar, B. V. et al. Human Tissue-Resident Memory T Cells Are Defined by Core Transcriptional and Functional Signatures in Lymphoid and Mucosal Sites. Cell Rep 20, 2921–2934, doi:10.1016/j.celrep.2017.08.078 (2017).

54. Mackay, L. K. et al. Hobit and Blimp1 instruct a universal transcriptional program of tissue residency in lymphocytes. Science 352, 459–463, doi:10.1126/science.aad2035 (2016).

55. Zundler, S. et al. Hobit-and Blimp-1-driven CD4(+) tissue-resident memory T cells control chronic intestinal inflammation. Nat Immunol 20, 288–300, doi:10.1038/s41590-018-0298-5 (2019).

56. Korn, T., Bettelli, E., Oukka, M. & Kuchroo, V. K. IL-17 and Th17 Cells. Annu Rev Immunol 27, 485–517, doi:10.1146/annurev.immunol.021908.132710 (2009).

57. Nowak, A. et al. CD137+CD154-Expression As a Regulatory T Cell (Treg)-Specific Activation Signature for Identification and Sorting of Stable Human Tregs from In Vitro Expansion Cultures. Front Immunol 9, 199, doi:10.3389/fimmu.2018.00199 (2018).

58. Brown, D. M., Kamperschroer, C., Dilzer, A. M., Roberts, D. M. & Swain, S. L. IL-2 and antigen dose differentially regulate perforin-and FasL-mediated cytolytic activity in antigen specific CD4+ T cells. Cell Immunol 257, 69–79, doi:10.1016/j.cellimm.2009.03.002 (2009).

59. Takeuchi, A. et al. CRTAM determines the CD4+ cytotoxic T lymphocyte lineage. J Exp Med 213, 123–138, doi:10.1084/jem.20150519 (2016).

60. White, A. M. & Wraith, D. C. Tr1-Like T Cells - An Enigmatic Regulatory T Cell Lineage. Front Immunol 7, 355, doi:10.3389/fimmu.2016.00355 (2016).

61. Hammond, S. A., Issel, C. J. & Montelaro, R. C. General method for the detection and *in vitro* expansion of equine cytolytic T lymphocytes. J Immunol Methods 213, 73–85, doi:10.1016/s0022-1759(98)00024-6 (1998).

62. Gruarin, P. et al. Eomesodermin controls a unique differentiation program in human IL-10 and IFN-gamma coproducing regulatory T cells. Eur J Immunol 49, 96–111, doi:10.1002/eji.201847722 (2019).

63. Zhang, P. et al. Eomesodermin promotes the development of type 1 regulatory T (TR1) cells. Sci Immunol 2, doi:10.1126/sciimmunol.aah7152 (2017).

64. Gregori, S., Goudy, K. S. & Roncarolo, M. G. The cellular and molecular mechanisms of immuno-suppression by human type 1 regulatory T cells. Front Immunol 3, 30, doi:10.3389/fimmu.2012.00030 (2012).

65. Yang, Y. H. et al. Deficiency of annexin A1 in CD4+ T cells exacerbates T cell-dependent inflammation. J Immunol 190, 997–1007, doi:10.4049/jimmunol.1202236 (2013).

66. D’Acquisto, F. et al. Annexin-1 modulates T-cell activation and differentiation. Blood 109, 1095–1102, doi:10.1182/blood-2006-05-022798 (2007).

67. D’Acquisto, F. et al. Impaired T cell activation and increased Th2 lineage commitment in Annexin-1-deficient T cells. Eur J Immunol 37, 3131–3142, doi:10.1002/eji.200636792 (2007).

68. Baaten, B. J. et al. CD44 regulates survival and memory development in Th1 cells. Immunity 32, 104–115, doi:10.1016/j.immuni.2009.10.011 (2010).

69. Shaffer, M. H. et al. Ezrin and moesin function together to promote T cell activation. J Immunol 182, 1021–1032, doi:10.4049/jimmunol.182.2.1021 (2009).

70. Scholzen, T. & Gerdes, J. The Ki-67 protein: from the known and the unknown. J Cell Physiol 182, 311–322, doi:10.1002/(SICI)1097-4652(200003)182:3<311::AID-JCP1>3.0.CO;2-9 (2000).

71. Sobecki, M. et al. Cell-Cycle Regulation Accounts for Variability in Ki-67 Expression Levels. Cancer Res 77, 2722–2734, doi:10.1158/0008-5472.CAN-16-0707 (2017).

72. Gerdes, J., Schwab, U., Lemke, H. & Stein, H. Production of a mouse monoclonal antibody reactive with a human nuclear antigen associated with cell proliferation. Int J Cancer 31, 13–20, doi:10.1002/ijc.2910310104 (1983).

73. Aleem, E., Kiyokawa, H. & Kaldis, P. Cdc2-cyclin E complexes regulate the G1/S phase transition. Nat Cell Biol 7, 831–836, doi:10.1038/ncb1284 (2005).

74. Diril, M. K. et al. Cyclin-dependent kinase 1 (Cdk1) is essential for cell division and suppression of DNA re-replication but not for liver regeneration. Proc Natl Acad Sci U S A 109, 3826–3831, doi:10.1073/pnas.1115201109 (2012).

75. Trapnell, C. et al. The dynamics and regulators of cell fate decisions are revealed by pseudotemporal ordering of single cells. Nat Biotechnol 32, 381–386, doi:10.1038/nbt.2859 (2014).

76. Alfei, F. et al. TOX reinforces the phenotype and longevity of exhausted T cells in chronic viral infection. Nature 571, 265–269, doi:10.1038/s41586-019-1326-9 (2019).

77. Khan, O. et al. TOX transcriptionally and epigenetically programs CD8(+) T cell exhaustion. Nature 571, 211–218, doi:10.1038/s41586-019-1325-x (2019).

78. Scott, A. C. et al. TOX is a critical regulator of tumour-specific T cell differentiation. Nature 571, 270–274, doi:10.1038/s41586-019-1324-y 10.1038/s41586-019-1324-y [pii] (2019).

79. Pearce, E. L. et al. Control of effector CD8+ T cell function by the transcription factor Eomesodermin. Science 302, 1041–1043, doi:10.1126/science.1090148 (2003).

80. Mazzoni, A. et al. Eomes controls the development of Th17-derived (non-classic) Th1 cells during chronic inflammation. Eur J Immunol 49, 79–95, doi:10.1002/eji.201847677 (2019).

81. Christophersen, A. et al. Distinct phenotype of CD4(+) T cells driving celiac disease identified in multiple autoimmune conditions. Nat Med 25, 734–737, doi:10.1038/s41591-019-0403-9 (2019).

82. Takeuchi, A. & Saito, T. CD4 CTL, a Cytotoxic Subset of CD4(+) T Cells, Their Differentiation and Function. Front Immunol 8, 194, doi:10.3389/fimmu.2017.00194 (2017).

83. Locafaro, G. et al. IL-10-Engineered Human CD4(+) Tr1 Cells Eliminate Myeloid Leukemia in an HLA Class I-Dependent Mechanism. Mol Ther 25, 2254–2269, doi:10.1016/j.ymthe.2017.06.029 (2017).

84. Gandhi, R. et al. Activation of the aryl hydrocarbon receptor induces human type 1 regulatory T cell-like and Foxp3(+) regulatory T cells. Nat Immunol 11, 846–853, doi:10.1038/ni.1915 (2010).

85. Grossman, W. J. et al. Differential expression of granzymes A and B in human cytotoxic lymphocyte subsets and T regulatory cells. Blood 104, 2840–2848, doi:10.1182/blood-2004-03-0859 (2004).

86. Culemann, S. et al. Locally renewing resident synovial macrophages provide a protective barrier for the joint. Nature 572, 670–675, doi:10.1038/s41586-019-1471-1 (2019).

87. Roncarolo, M. G., Gregori, S., Bacchetta, R. & Battaglia, M. Tr1 cells and the counter-regulation of immunity: natural mechanisms and therapeutic applications. Curr Top Microbiol Immunol 380, 39–68, doi:10.1007/978-3-662-43492-5_3 (2014).

88. Zimmermann, J. et al. T-bet expression by Th cells promotes type 1 inflammation but is dispensable for colitis. Mucosal Immunol 9, 1487–1499, doi:10.1038/mi.2016.5 (2016).

89. Jain, A. et al. T cells instruct myeloid cells to produce inflammasome-independent IL-1beta and cause autoimmunity. Nat Immunol 21, 69, doi:10.1038/s41590-019-0559-y (2020).

90. Hamilton, J. A. GM-CSF in inflammation. J Exp Med 217, doi:jem.20190945 [pii] 10.1084/jem.20190945 (2020).

91. Gunn, M. D. et al. A B-cell-homing chemokine made in lymphoid follicles activates Burkitt’s lymphoma receptor-1. Nature 391, 799–803, doi:10.1038/35876 (1998).

92. Legler, D. F. et al. B cell-attracting chemokine 1, a human CXC chemokine expressed in lymphoid tissues, selectively attracts B lymphocytes via BLR1/CXCR5. J Exp Med 187, 655–660, doi:10.1084/jem.187.4.655 (1998).

93. Zotos, D. et al. IL-21 regulates germinal center B cell differentiation and proliferation through a B cell-intrinsic mechanism. J Exp Med 207, 365–378, doi:10.1084/jem.20091777 (2010).

94. Spolski, R. & Leonard, W. J. Interleukin-21: basic biology and implications for cancer and autoimmunity. Annu Rev Immunol 26, 57–79, doi:10.1146/annurev.immunol.26.021607.090316 (2008).

95. Rao, D. A. T Cells That Help B Cells in Chronically Inflamed Tissues. Front Immunol 9, 1924, doi:10.3389/fimmu.2018.01924 (2018).

96. Sedger, L. M. & McDermott, M. F. TNF and TNF-receptors: From mediators of cell death and inflammation to therapeutic giants - past, present and future. Cytokine Growth Factor Rev 25, 453–472, doi:10.1016/j.cytogfr.2014.07.016 (2014).

97. Hradilkova, K. et al. Regulation of Fatty Acid Oxidation by Twist 1 in the Metabolic Adaptation of T Helper Lymphocytes to Chronic Inflammation. Arthritis Rheumatol 71, 1756–1765, doi:10.1002/art.40939 (2019).

98. Lin, C. C. et al. Bhlhe40 controls cytokine production by T cells and is essential for pathogenicity in autoimmune neuroinflammation. Nat Commun 5, 3551, doi:10.1038/ncomms4551 (2014).

99. Huynh, J. P. et al. Bhlhe40 is an essential repressor of IL-10 during Mycobacterium tuberculosis infection. J Exp Med 215, 1823–1838, doi:10.1084/jem.20171704 (2018).

100. Martinez-Llordella, M. et al. CD28-inducible transcription factor DEC1 is required for efficient autoreactive CD4+ T cell response. J Exp Med 210, 1603–1619, doi:10.1084/jem.20122387 (2013).

101. Stuart, T. et al. Comprehensive Integration of Single-Cell Data. Cell 177, 1888–1902 e1821, doi:10.1016/j.cell.2019.05.031 (2019).

102. Weiner, J. & Domaszewska, T. tmod: an R package for general and multivariate enrichment analysis. doi:10.7287/PEERJ.PREPRINTS.2420V1 (2016).

103. Yamaguchi, K. D. et al. IFN-beta-regulated genes show abnormal expression in therapy-naive relapsing-remitting MS mononuclear cells: gene expression analysis employing all reported protein-protein interactions. J Neuroimmunol 195, 116–120, doi:10.1016/j.jneuroim.2007.12.007 (2008).

104. Zyla, J. et al. Gene set enrichment for reproducible science: comparison of CERNO and eight other algorithms. Bioinformatics, doi:10.1093/bioinformatics/btz447 (2019).

